# Phosphatidylinositol 3-phosphate mediates Arc capsids secretion through the multivesicular body pathway

**DOI:** 10.1101/2023.12.19.572392

**Authors:** Kritika Mehta, Henry Yentsch, Jungbin Lee, Tianyu Terry Gao, Kai Zhang

## Abstract

Activity-regulated cytoskeleton-associated protein (Arc/Arg3.1) is an immediate early gene that plays a vital role in learning and memory. The recent discovery that Arc mediates the inter-neuronal RNA transfer implies its role in regulating neuronal functions across long distances. Arc protein has structural and functional properties similar to viral Group-specific antigen (Gag). By assembling into high-order, virus-like capsids, Arc mediates the intercellular RNA transfer. However, the exact secretion pathway through which Arc capsids maneuver cargos is unclear. Here, we identified that Arc capsids assemble and secrete through the endosomal-multivesicular body (MVB) pathway. Arc’s endosomal entry is likely mediated by phosphatidylinositol-3-phosphate (PI3P). Indeed, reconstituted Arc protein preferably binds to PI3P. In mammalian cells, Arc forms puncta that colocalizes with FYVE, an endosomal PI3P marker, and competitive binding to PI3P via prolonged FYVE expression reduces the average number of Arc puncta per cell. Overexpression of MTMR1, a PI3P phosphatase, significantly reduces Arc capsid secretion. Arc capsids secrete through the endosomal-MVB axis as extracellular vesicles. Live-cell imaging shows that fluorescently labeled Arc primarily colocalizes Rab5 and CD63, early endosomal and MVB markers, respectively. Superresolution imaging resolves Arc accumulates within the intraluminal vesicles of MVB. CRISPR double knockout of RalA and RalB, crucial GTPases for MVB biogenesis and exocytosis, severely reduces Arc-mediated RNA transfer efficiency. These results suggest that, unlike the Human Immunodeficiency Virus Gag, which assembles on and bud off from the plasma membrane, Arc capsids assemble at the endocytic membranes of the endosomal-MVB pathway mediated by PI3P. Understanding Arc’s secretion pathway helps gain insights into its role in intercellular cargo transfer and highlights the commonality and distinction of trafficking mechanisms between structurally resembled capsid proteins.

## Introduction

On average, each neuron cell forms approximately 1000 synapses, proximal connections between cells that facilitate information relay across cells mediated by neurotransmitters. Neuronal activity at a synapse could stimulate a long-distance synapse-to-nucleus communication to activate the transcription of immediate early genes in the stimulated cells. One such actively transcribed gene is Activity Regulated Cytoskeleton-associated protein (Arc/Arg3.1), whose mRNA is transported to neuronal dendrites in response to synaptic activity (1–3). The local translation of Arc mRNA produces Arc protein, which mediates synaptic protein trafficking and modifies local proteome, leading to morphological and functional changes of synaptic strength between associated cells. Such a “fire-together, wire-together” Hebbian synaptic plasticity is believed to underpin molecular mechanisms underlying learning, memory formation, and long-term memory consolidation.

While synapse-to-nucleus communication represents a unique mode of action for long-distance communication in polarized neuronal cells, other mechanisms exist in nature to facilitate proximity-independent communication between mammalian cells (4–19). For example, retroviruses package their genome into capsids, a supramolecular structure that could undergo exocytosis from a parent cell, diffuse through the extracellular space, and infect a recipient cell. Such long-distance delivery of capsids between cells necessitates infrastructure for stability and sufficient volume. Extracellular vesicles (EVs) are among such infrastructures that play an effective role in intercellular communications (20–26). The spatial and temporal release of EVs into the extracellular space is regulated by the cellular machinery involved in its biogenesis and exocytosis (22). Exosomes are a class of EVs that employ the multivesicular body (MVB) pathway to generate membrane-bound EVs inside the cell’s cytoplasm (27). Microvesicles, on the other hand, release via outward budding of the plasma membrane (28).

It was not expected that such a capsid-mediated pathway could have implications in neuronal communication until recently when Arc was found to regulate intercellular RNA transfer. In this process, Arc proteins assemble into high-order, virus-like capsids encapsulating its RNA, exocytose from the parent cell, and infect a remote recipient neuron (14, 15). It is not completely surprising for Arc’s “new” role, considering Arc is a retrotransposon-derived gene, but structurally resembled protein could evolve distinct trafficking pathways with unique regulators or mediators (29–31). Phospholipids play a crucial role in retrovirus capsid assembly and trafficking. For example, stable attachment of the Human Immunodeficiency virus (HIV) Gag to the membrane requires its specific interaction with phosphatidylinositol-4,5-bisphosphate [PI(4,5)P2] enriched at the plasma membrane (32, 33). However, it is unclear if phospholipids regulate Arc capsid biogenesis and trafficking (34).

Here, we employ a combination of genetic, imaging, and biochemical analysis to understand the biogenesis and trafficking of Arc-containing capsids. Unlike HIV particles, which assemble proximal to the plasma membrane, Arc capsid assembly and secretion proceed through the endosomal-MVB pathway. Intriguingly, the high affinity of Arc to PI3P, validated both in reconstituted Arc protein and in mammalian cells, mediates Arc capsid trafficking. Competitive binding or modulation of PI3P levels in live cells through over-expression of MTMR1, a PI3P phosphatase, significantly reduces Arc-containing EV secretion. Fluorescently labeled Arc colocalizes with endosomal and MVB markers in mammalian cells. Superresolution imaging resolves oligomeric Arc within the intraluminal vesicles of MVB. Genetic knockout of RalA and RalB, crucial GTPases for MVB biogenesis and exocytosis, severely reduces Arc-mediated RNA transfer efficiency. These results suggest that Arc utilizes the endosomal-MVB axis for capsid assembly and secretion. Understanding Arc capsid assembly and trafficking will shed light on the commonality and distinction of molecular machinery regulating structurally resembled capsid proteins. Knowledge of Arc intercellular trafficking could provide a better understanding of brain physiology and pathology, as it has been found that various Arc mutants occur in patients with neurological and neuropsychiatric diseases such as Alzheimer’s disease, autism spectrum disorder, and Schizophrenia (35, 36).

## Results

### HaloArc mediates intercellular RNA transfer and protein expression

To visualize the Arc protein dynamics in live cells, we constructed HaloArc, a plasmid containing mouse Arc with an N-terminal fusion of HaloTag, flanked by the 5’-UTR and 3’-UTR of the Arc mRNA (**Fig. S1A**). UTRs were included because of their potential function in mediating intercellular RNA transfer (14). We first confirmed that this synthetic Arc maintains the capacity of intercellular RNA transfer in HEK293T cells, which express no endogenous Arc but allow for validation of Arc RNA transfer shown in previous work (14). We developed the donor-recipient assay as depicted in **Fig. 1A**. HaloArc was transfected in HEK293T cells (donor cells), expression was confirmed by fluorescence microscopy, and their conditioned medium was collected and centrifuged. The supernatant was administrated to a fresh batch of non-transfected HEK293T cells (recipient cells). After incubating the conditioned medium with the recipient cells for 24 h, the expression of HaloArc was measured by fluorescence microscopy. In parallel, extracellular vesicles (EV) were extracted from part of the conditioned medium for further analysis. A successful HaloArc-mediated donor-recipient RNA transfer assay fulfills the following criteria: 1) upregulated HaloArc protein expression in the recipient cells, 2) enrichment of HaloArc protein in the EVs, and 3) enrichment of HaloArc RNA in the EVs.

**Figure 1.**
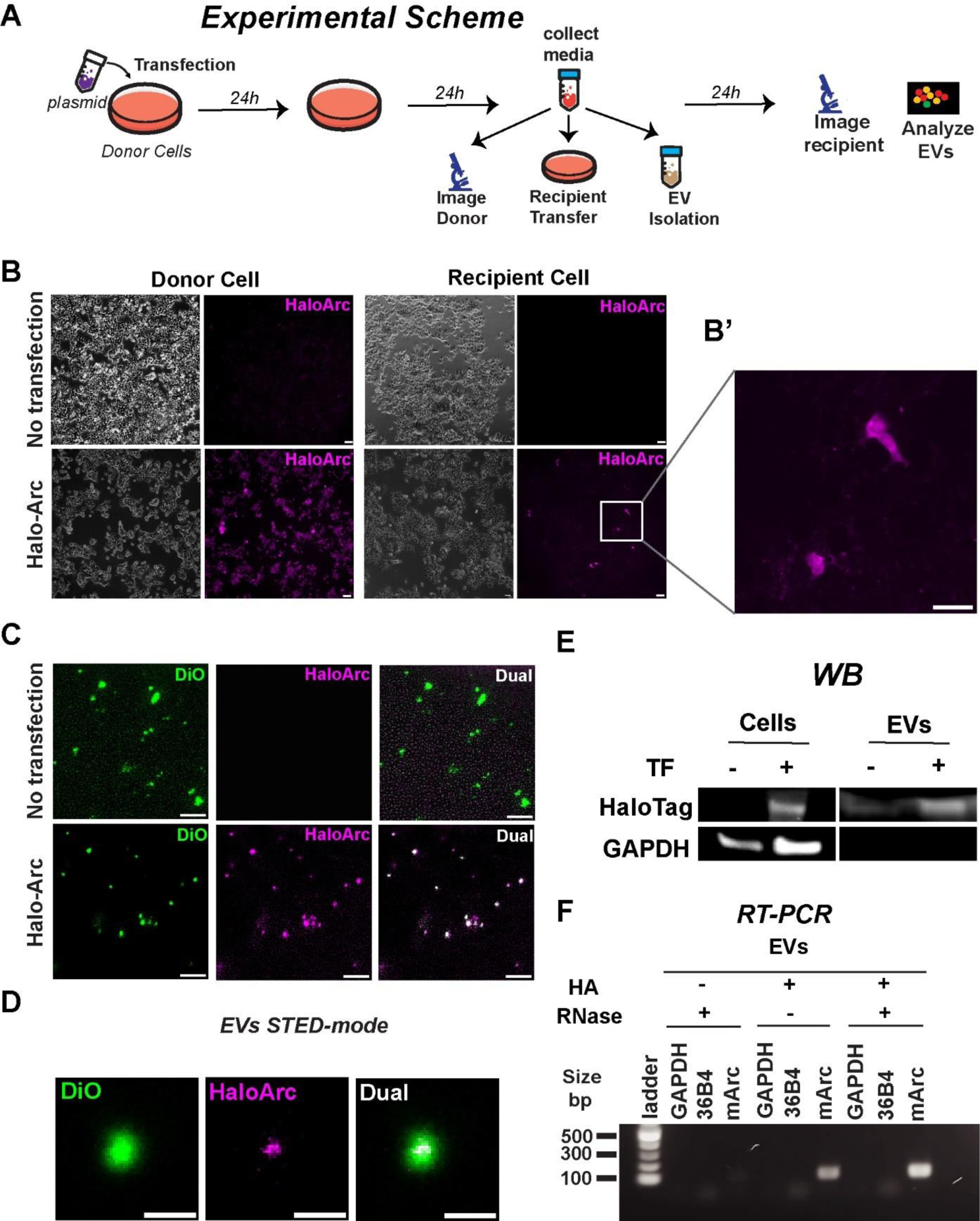
Halo-Arc can transfer between HEK293T cells. A) Experimental scheme for analyzing HaloArc function: Media transfer from HaloArc transfected cells was treated to the untransfected recipient cells to confirm material transfer. Extracellular vesicles (EVs) were also purified from the transfected cell media for further downstream analysis. B) Donor-recipient assay showing HA fluorescence with JF549 Halo ligand staining in the donor cells (left) and recipient cells (right), where untransfected donor cells are negative control. B’) Inset shows the recipient cells expressing HaloArc after media treatment. Scale bar: 50 µm. C) Epifluorescence imaging of EVs isolated from untransfected and HaloArc transfected cell media and labeled with 5 μM DiO (green) and 100nM JF549 Halo ligand (magenta). Scale bar: 2 µm. D) Representative 2D-STED image of an EV stained with DiO (green) and JF-646 halo ligand (magenta) capturing HaloArc resides inside the EVs. Scale bar: 500 nm. E) Western blot showing expression of HaloArc in the transfected cells and their EVs isolated from the conditioned media (no transfection serves as negative control F) Reverse transcription PCR showing expression of HaloArc mRNA inside the EV fraction with and without RNase treatment.

Expression of HaloArc in the donor cells was confirmed by Western blot with a primary antibody against HaloTag. Compared to non-transfection control, overexpression of HaloArc in HEK293T cells resulted in a single band at the expected molecular weight of 75 kD within the whole cell lysate (**Fig. S1B**). Expression of HaloArc was also confirmed by fluorescence imaging, where JF549-labeled HaloTag ligand was used to label HaloTag in live cells. In donor HEK293T cells transfected with HaloArc, more than 80% of cells showed strong fluorescence from JF549. Non-transfected cells showed no detectable fluorescence (**Fig. 1B, left)**. When the same labeling procedure was applied to recipient cells incubated with the conditioned medium of transfected donor cells, a similar enhancement of fluorescence was found, indicating successful expression of HaloArc in the recipient cells (criterion #1) (**Fig. 1B, right, Fig. 1B’**). We further confirmed that Arc-containing EVs were membrane-bound by co-staining Arc with DiO, a lipophilic tracer with green fluorescence. Arc-positive EVs unanimously showed green fluorescence, indicating the presence of the membrane structure (**Fig. 1C**). The average fluorescence intensity of Arc-expressing recipient cells was slightly less than that of Arc-expressing donor cells. However, both were significantly above the basal level from the non-Arc-expressing cells in the same culture (**Fig S1C**). To determine the relative location of the Arc protein with respect to the EV membrane, we applied two-color Stimulated Emission Depletion (STED) super-resolution imaging to resolve both the membrane and Arc proteins. Arc protein localized within the membrane structure, consistent with a membrane-bound exosome encapsulating Arc capsid (**Fig. 1D**). The full-width half-maximum calculated from the one-dimensional Gaussian fit to a single EV in STED-mode provided us with an average diameter of 303 ± 26 nm (n=5 single EVs imaged in STED-mode, **Fig.S1D**). Western blot analysis of purified EVs reveals the presence of HaloArc, meeting criterion #2 (**Fig. 1E**). Finally, RT-PCR measurement reveals that mArc RNA resides in EVs, likely within the capsids, because RNase treatment did not degrade the RNA, fulfilling criterion #3 (**Fig. 1F**). We noted that the band intensity of the RNase(+) sample is higher than that of the RNase(-) sample, likely because RNase treatment enriched HaloArc RNA by degrading other EV-residing RNAs outside the capsids. These results confirmed that HaloArc functions similarly to endogenous Arc in mediating the intercellular transfer of RNA, which can be expressed in the recipient cells.

### HaloArc confines into slow-diffusing, membrane-associated puncta in mammalian cells

We reasoned that snapshots of the subcellular distribution and morphology of HaloArc could imply their protein trafficking pathways. A home-built highly inclined and laminated optical sheet (HILO) microscopy was used for single-molecule imaging inside cells (37). When expressed and fluorescently labeled in HEK293T cells, HaloArc formed slow-diffusing, almost static puncta in the cytoplasm (**Fig. 2A(i-iii), Movie S1**). This structure was not an artifact of HaloTag because fluorescently labeled HaloTag alone showed typical fast diffusion in HEK293T cells (**Fig. 2B(i-iii), Movie S2**). We collected diffusion trajectories from 1770 HaloTag molecules and 624 HaloArc puncta and constructed a histogram of their diffusion coefficients (**Fig. 2A(iv),2B(iv)**). The probability distribution of both histograms appeared to be lognormal with similar widths. The mean diffusion coefficient of HaloTag alone is 1.0 μm^2^/s, almost two orders of magnitude larger than that of HaloArc (0.01 μm^2^/s) (**Fig. 2C**). The HaloArc puncta are significantly brighter than HaloTag alone, indicating the clustering of Arc molecules. HaloArc puncta are associated with the membrane because DiO-positive puncta colocalized with HaloArc (**Fig. 2D**). When HaloTag was replaced by superfolder GFP (sfGFP), sfGFP-Arc also formed large puncta in HEK293T cells, indicating that puncta formation is not likely caused by fusion proteins (**Fig. 2E**). We further confirmed that HaloArc forms puncta in various mammalian cell lines, such as the neuroblastoma SH-SY5Y (**Fig. 2F**) and the fibroblast NIH3T3 (**Fig. 2G**), as well as the primary rat cortical neuronal cultures transduced by lentivirus encoding HaloArc show similar puncta (**Fig. 2H, Fig. S1E**).

**Figure 2.**
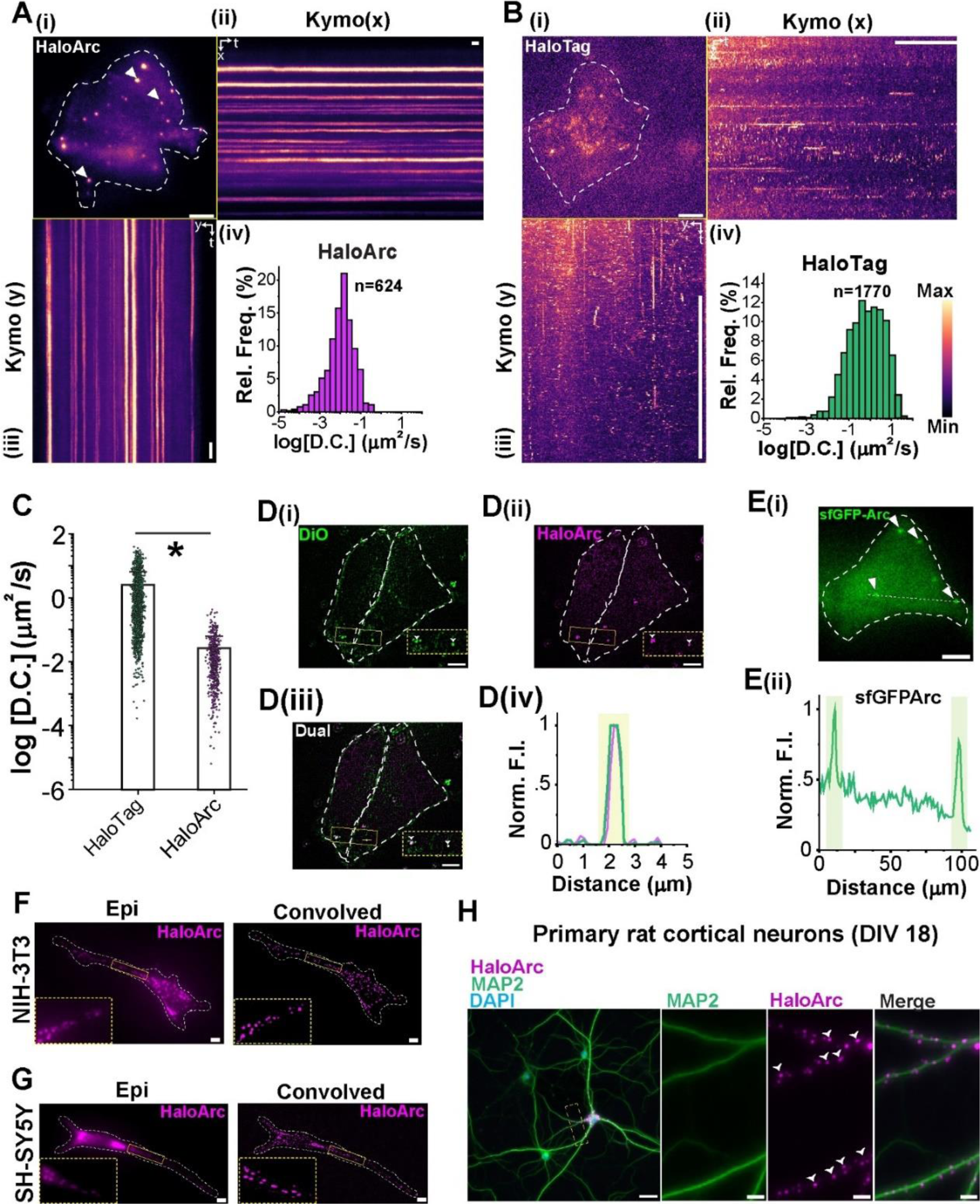
Halo-Arc assembles into membrane-associated clusters in live cells. A(i) Representative HILO image of a HEK293T cell expressing HaloArc stained in JF549 halo ligand (arrows indicate the HaloArc clusters). Scale bar: 5 µm. A(ii-iii) X and Y kymograph showing the horizontal and vertical trajectory across the entire cell; scale is 1s. A(iv) Histogram for Diffusion coefficient for tracked halo-Arc particles (n=624 particles; N=3 experiments). B(i) Representative HILO image of a HEK293T cell expressing halotag and stained in JF549 halo ligand. Scale bar: 5 µm. B(ii-iii) X and Y kymograph showing horizontal and vertical trajectory across the entire cell; scalebar: 1 s. B(iv) Histogram for diffusion coefficients for tracked halotag particles (n=1770 particles; N=3 experiments). C) Scatter plot of the diffusion coefficients of HaloArc and halotag particles tracked (Mann-Whitney test; *: p <<0.05). D(i-ii) Representative convolved epifluorescence image of transfected HEK293T cells stained with DiO (left, green) co-stained with JF549 halo for HaloArc (right, magenta). D(iii) Merged image of D(i-ii) in both the channels (arrows indicate HaloArc clusters localized within DiO-stained vesicles. D(iv) Intensity profile across the ROI in D(iii). Scale bar: 5 µm. E(i) Representative epifluorescence image of a HEK293T cell expressing sfGFP-Arc (arrows indicate Arc clusters). Scale bar: 5 µm. E(ii) Intensity profile across the ROI in E(i) with peaks indicating the large Arc clusters, as highlighted. F) Representative epifluorescence (left) and convolved (right) image of a stained SH-SY5Y cell expressing HaloArc showing the cluster phenotype. Inset shows the ROI marked to show the clusters inside the cell. Scale bar: 5 µm. G) Representative epifluorescence (left) and convolved (right) image of a stained NIH-3T3 cell expressing HaloArc showing the cluster phenotype. Inset shows the ROI marked to show the clusters inside the cell. Scale bar: 5 µm. H) Representative epifluorescence image of HaloArc transduced and stained primary rat cortical neurons showing MAP2 (green), nucleus (blue) and HaloArc (magenta). Scale bar in H(left): 20 µm. Enlarged images of the ROI marked (left) show the HaloArc clusters inside neuronal processes. Scale bar: 5 µm.

### Recombinant Arc proteins assemble into capsids

The slow-diffusing and bright Arc puncta indicate the clustering of Arc molecules inside cells. To further understand Arc assembly, we purified Arc from bacteria using GST affinity chromatography (**Figure 3A, S2A**). Arc protein was also run on a size exclusion column with the TBS buffer (pH 7.4) and was eluted into three peaks with the estimated sizes of 670 kDa, 200 kDa, and 100 kDa (**Fig. S2B**). All three fractions were pooled together and used for protein characterization (**Fig. S2C**). Western blot analysis with purified Arc protein showed a single band at approximately 50 kDa probed by an Arc primary antibody (**Fig. 3B**). Mass spectroscopy further confirms the identity of purified Arc (**Fig. S2D-E**). Consistent with previous work (14), incubation with 500 mM sodium phosphate buffer shifted the size of Arc from 14 nm to 30 nm in dimeter as measured by dynamic light scattering (DLS) (**Fig. 3C**). Arc protein treated with phosphate and mixed with RNA in 10:1 (w/w) ratio was also imaged using negative staining Transmission Electron Microscopy (TEM), which clearly revealed that Arc formed capsid structure with an average diameter of 32.9 nm (n=201 particles), consistent with DLS measurement (**Fig. 3D-F**). Capsid induction appeared to be independent of RNA sequences, as RNAs from whole cells, GFP, and Arc all induce capsid formation in the high phosphate buffer (**Fig. S2F**). These results confirmed that purified Arc protein can assemble into capsid-like structures.

**Figure 3.**
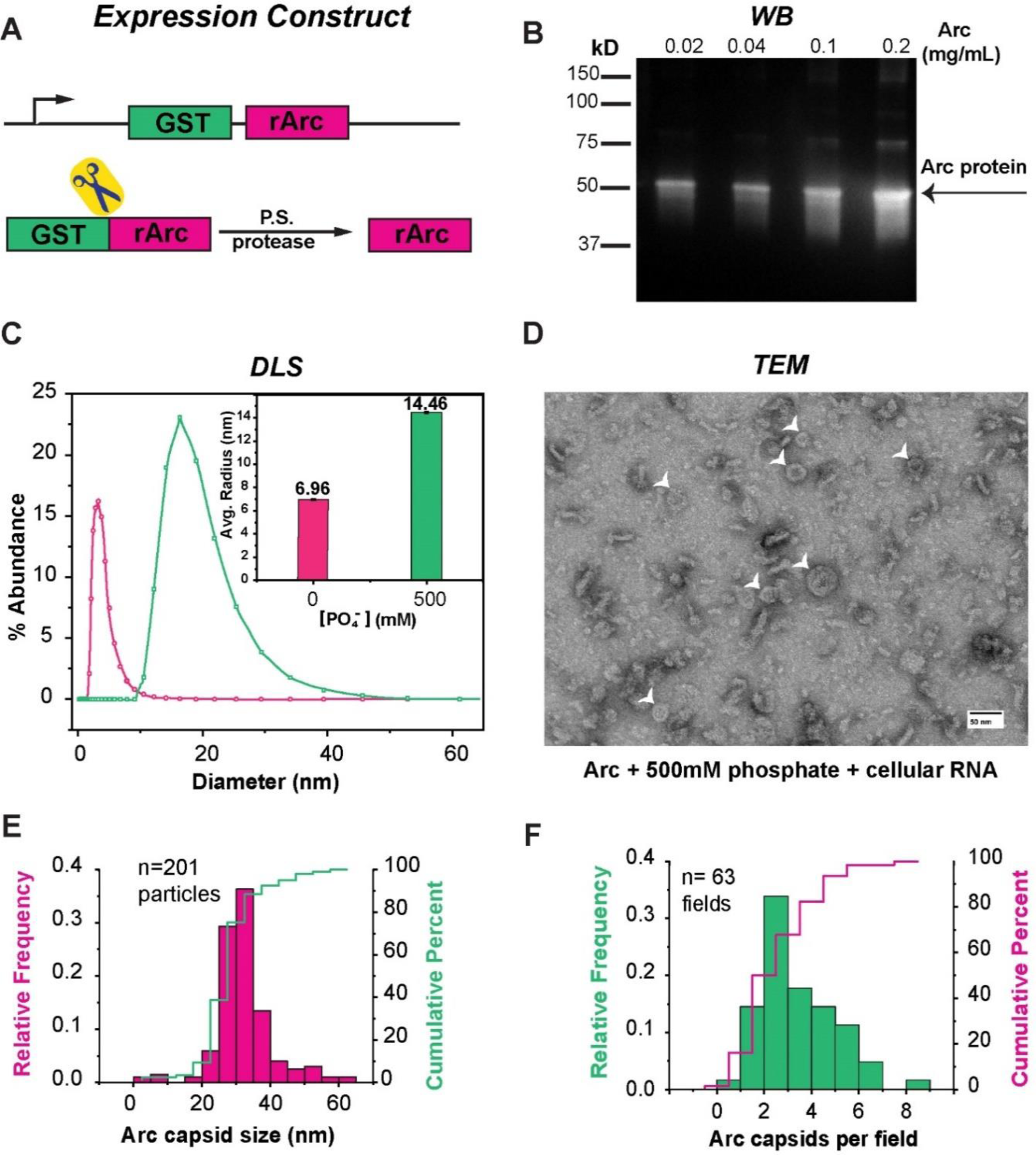
Purified Arc protein can assemble into capsids. A) Schematic of the GST-Arc construct used for affinity purification of Arc protein after on-column cleavage of the GST tag using the PreScission (P.S.) protease. B) Arc protein identity confirmation using western blot against the Arc antibody against different concentrations of purified Arc protein. C) DLS analysis showing Arc protein size with (green) and without (pink) high phosphate buffer treatment, 500 mM sodium phosphate (inset shows the average radius measured from DLS for both the conditions). D) TEM micrograph of Arc capsids induced with RNA and phosphate buffer treatment of the purified Arc protein (arrows indicate the typical double-shelled capsids of Arc). E) Quantification of capsid diameter induced with purified Arc protein (n=201 capsids; N=4 experiments). F) Quantification of the number of Arc capsids per TEM field (n=63 fields; N=4 experiments).

### Arc proteins are associated preferably with PI3P

Phospholipids play a crucial role in regulating protein trafficking in live cells. Distinct pools of phospholipids reside in the membranes of different intracellular vesicles. We reason that Arc assembly inside cells could be mediated through phospholipids. To determine whether Arc selectively binds to phospholipids, we performed a lipid-protein interaction assay by incubating purified Arc protein with a membrane lipid strip, a membrane pre-spotted with 15 different types of lipids. Intriguingly, Arc proteins are preferably associated with monophosphorylated phosphatidylinositol, phosphatidylinositol-3-phosphate (PI3P), phosphatidylinositol-4-phosphate (PI4P), and phosphatidylinositol-3-phosphate (PI5P) (**Fig. 4A**). In particular, PI3P is associated with Arc with approximately 2-fold and 3-fold stronger affinity than PI4P and PI5P, respectively (**Fig. 4B**).

**Figure 4.**
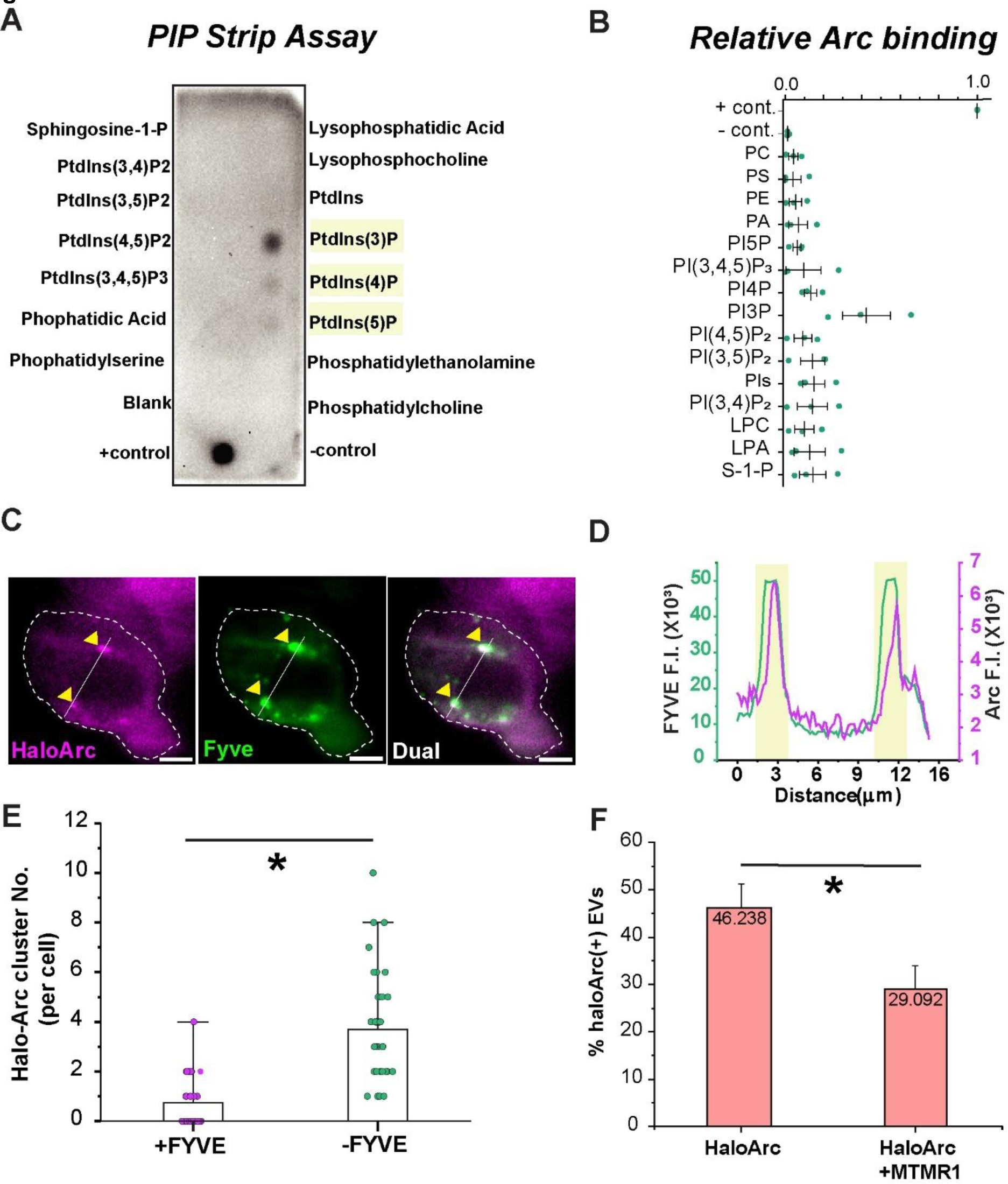
Arc protein interacts with PI3P. A) PIP-strip lipid-overlay assay showing preferable Arc binding to monophosphorylated phosphoinositides (highlighted in yellow). Arc protein is probed using the Arc antibody. B) Quantification of PIP-strip signal intensity highlighting Arc has the highest affinity towards PI3P. C) Representative 2-color HILO image of a HEK293T cell co-expressing 2×Fyve-GFP (green; a PI3P marker) and HaloArc (magenta). Arrows highlight HaloArc clusters colocalized with the Fyve domain (dual). Scale bar: 5 μm. D) Fluorescence intensity quantification of the ROI in C). Colocalized clusters are shown in yellow. E) Quantification of the count of HaloArc clusters per cell under the HILO field with (pink; n=34 cells) or without (green; n=35 cells) FYVE co-transfection. F) EV counting analysis of EVs from HaloArc and HaloArc+MTMR1 expressing cells. DiO staining was used as a normalization factor for counting. (Mann-Whitney test; *: p<0.05).

Motivated by this result, we tested if Arc binds to PI3P in cells. The FYVE finger (zinc finger originally observed in Fab1p, YOTB, Vac1p and EEA1) protein functions in the membrane recruitment of cytosolic proteins by binding to PI3P, which is found primarily on endosomes and is commonly used as a PI3P marker in cells (38). We co-transfected HaloArc with 2×FYVE-EGFP in HEK293T cells and performed two-color HILO imaging (**Fig. S3A, Movie S3**). Consistent with the PIP-strip result, we observed colocalization between HaloArc and 2×FYVE-EGFP (**Fig. 4C-D**), indicating Arc’s preferable binding to PI3P. Curiously, the number of Arc puncta (no FYVE, Arc clusters=3.7±2.3; n=34 cells) is reduced when co-transfected with 2×FYVE-EGFP (transfected with FYVE, Arc clusters= 0.7±1.0; n=35 cells), which implies that FYVE’s competitive binding to PI3P affects Arc puncta formation (**Fig. 4E**). Interestingly, a previous study on PIP-regulated gephyrin clustering also presented a similar reduction in gephyrin clusters with prolonged FYVE overexpression, possibly due to competitive binding of FYVE to membrane PI3P (39). To further understand the effect of PI3P on Arc exocytosis, we co-expressed MTMR1, a PI3P phosphatase, along with HaloArc. We then isolated the EVs and stained them with both DiO (for normalization) and halo staining. We observed that co-expression of MTMR1 significantly reduced HaloArc secretion (**Fig. 4F**). These results suggest that PI3P may mediate the endosomal entry of Arc protein inside cells.

### HaloArc clusters colocalize with markers for early endosome and MVB in mammalian cells

PI3P is enriched on the limiting membrane of early endosomes and intraluminal vesicles within MVBs (38); therefore, we proceeded to determine the identity of Arc-associated vesicles along the endosomal pathways. We carried out co-localization experiments between HaloArc and various markers of the endo-lysosomal system, including Rab5-EGFP (early endosomal), Rab7-EGFP (late endosomal and MVB), and sfGFP-CD63 (MVB). Because of the potential aberration of far-field optical microscopy, single snapshots of dual-color imaging may introduce artifacts about particle colocalization. If two distinct vesicles were located in a stacked position along the axial focus, they could be mistakenly assigned as colocalized particles. To address this issue, we used dual-color single-particle tracking to follow the trafficking trajectory of vesicles under both emission channels. When the puncta in both green (vesicle marker) and red (Arc) channels move together during the whole 200-second data acquisition time, markers for both channels are more likely to reside on the same puncta (**Fig. 5A-C, Movie S4A-C**). We found that Arc puncta colocalized with 58.3% Rab5+ (n=24), 81.0% CD63+ (n=89), and 86.7% Rab7+ vesicles (n=27) (**Fig. 5D**). We also imaged sfGFP-Arc with another MVB marker protein mCherry-TSG101 and found 94.73% colocalization (n=20 vesicles, **Fig. S4A-B, S4E, Movie S4D**). Colocalization studies with Rab7-EGFP and mCherry-TSG101 showed 93.3% colocalization of both markers (n= 30 vesicles, **Fig. S4C-E**, **Movie S4E**), indicating that both Rab7 and TSG101 are effective MVB markers.

**Figure 5.**
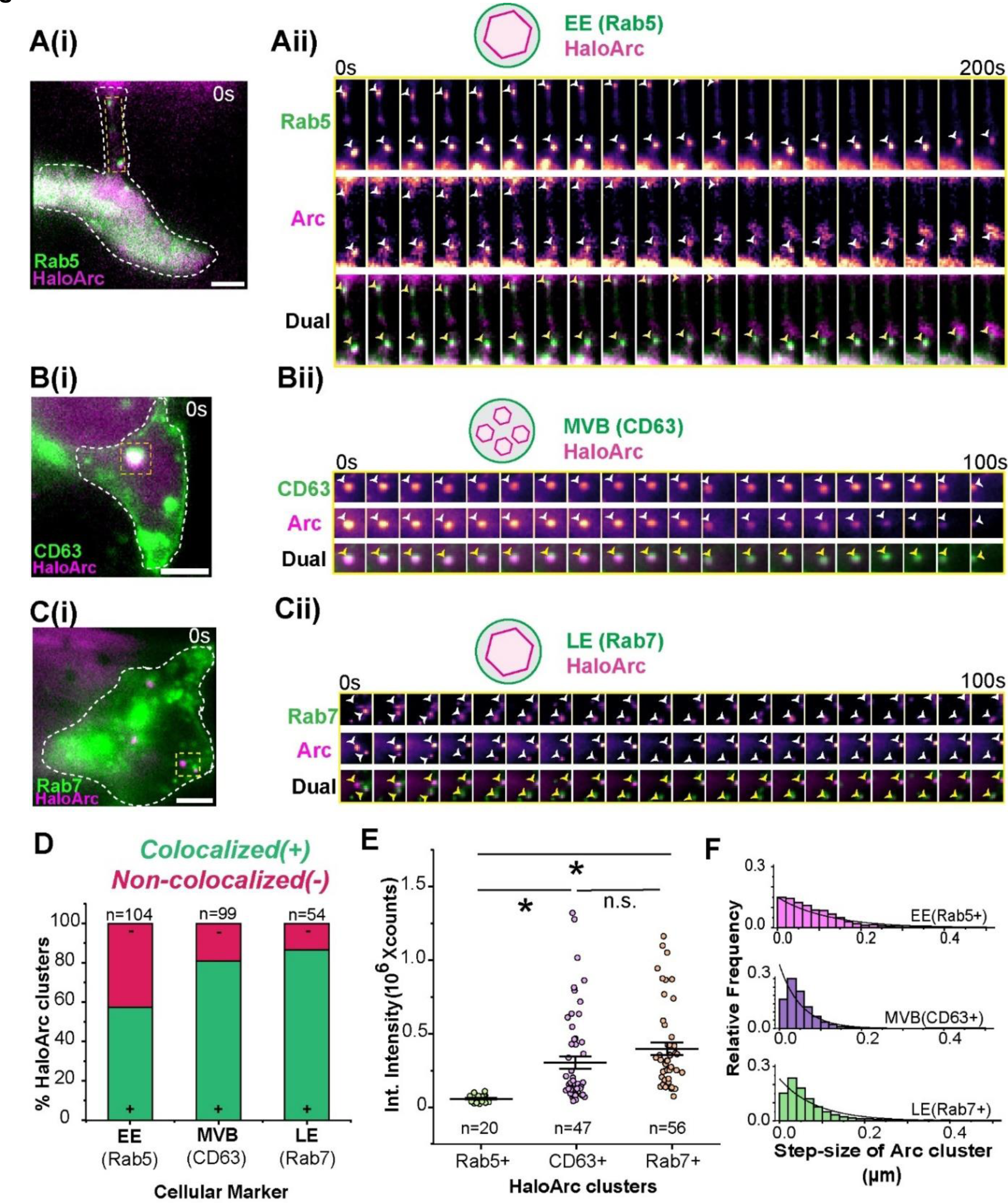
Arc clusters exist inside different endosomal species. A(i) A representative 2-color HILO image of HEK293T cells co-transfected with HaloArc (magenta) and Rab-GFP (green). A(ii) Montage of the same cell ROI marked in yellow in A(i) for Rab5 (top), HaloArc (middle) and merged Rab5 and HaloArc (bottom). Arrows indicate HaloArc colocalized within Rab5+ vesicles. B(i) A representative 2-color HILO image of HEK293T cells co-transfected with HaloArc (magenta) and CD63-sfGFP (green). B(ii) Montage of the same cell ROI marked in yellow in B(i) for CD63 (top), HaloArc (middle) and merged CD63 and HaloArc (bottom). Arrows indicate HaloArc colocalized within CD63+ vesicles. C(i) A representative 2-color HILO image of a HEK293T cell co-transfected with HaloArc (magenta) and Rab7-GFP (green). C(ii) Montage of the same cell ROI marked in yellow in C(i) for Rab7 (top), HaloArc (middle) and merged Rab7 and HaloArc (bottom). Arrows indicate HaloArc colocalized within Rab7+ vesicles. Warmer colors in single-channel montages imply higher intensities. D) Quantification for a fraction of HaloArc clusters colocalizing with each of the endosomal species (Rab5: early endosome, n=104 clusters; CD63: multivesicular body, n=99 clusters; and Rab7: late endosome, n=54 clusters) with green depicting vesicle(+) and red depicting vesicle(-) HaloArc cluster for each endosomal marker. E) Scatter plot of integrated fluorescence intensities of all the HaloArc clusters within an endosomal marker (Rab5, n=20; CD63, n=56; Rab7, n=47). F) Histogram of HaloArc cluster step size within each type of endosome. (Mann-Whitney test; *: p<0.05). Scale bar: 5 μm for all images.

Intensity analysis shows that early endosome-associated Arc puncta (colocalized with Rab5, n=20) showed a lower fluorescence intensity than those associated with late endosomes or MVBs (Rab7, n=47; CD63, n=56) (**Fig. 5E, S4F**). Analysis of HaloArc puncta step size (100 ms frame rate) shows that early endosome-associated HaloArc has a larger step size than those colocalized with MVBs and late endosomes (**Fig. 5F**). These results indicate that Arc proteins might either acquire different oligomeric states or progressively enriched as they proceed through the endosome-MVB pathway.

### RalA and RalB double knockout alters the morphology of HaloArc clusters in mammalian cells

Considering that purified Arc protein binds to PI3P and Arc puncta colocalizes with early endosomes, late endosomes, and MVB, and Arc punta intensity progressively increases from early endosomes to MVB, we hypothesize that small oligomeric Arc could enrich at the early endosomes, and assemble capsids as Arc enters MVB (likely within the intraluminal vesicles), which eventually undergoes membrane fusion at the plasma membrane to release extracellular vesicles (EV) containing Arc capsid. To test this hypothesis, we manipulated the MVB biogenesis and compared phenotypic changes of Arc puncta. Ral GTPase is crucial for both MVB biogenesis and exocytosis (27, 40, 41). We, therefore, deleted endogenous RalA and RalB, two isoforms of Ral GTPase by CRISPR-knockout in HEK293T cells. RalA and RalB double knockout (DKO) was confirmed using qRT-PCR in a polyclonal population upon selection of the cells treated with the CRISPR-Cas9 machinery (**Fig S5A**). We further selected a monoclonal colony that had no expression for RalA and RalB (double knockout or DKO) mRNA for further analysis (**Fig. S5B**).

Similar to wild-type HEK293T cells, HaloArc forms puncta in DKO cells (**Fig. 6A-B**). These puncta colocalize with FYVE as in WT cells (**Fig. S5C**). However, DKO cells show more Arc puncta per cell than wild-type cells (**Fig. 6C**, n= 9 cells for WT, n= 35 cells for DKO). In addition, the fluorescence intensity per puncta in DKO cells appears higher, with a larger average and integrated intensity than that of wild-type cells (**Fig. 6D, S5D-E,** n=87 clusters for WT, n= 411 clusters for DKO). Additionally, puncta in DKO appeared to be larger (**Fig. S5F**) and less circular (**Fig. S5G**) than those in WT cells. These differences in puncta intensity and morphology imply erroneous MVB biogenesis in the DKO cells. In support of this idea, CD63-EGFP (MVB marker) positive vesicles were larger in DKO cells than those in wild-type cells (**Fig. 6E**).

**Figure 6.**
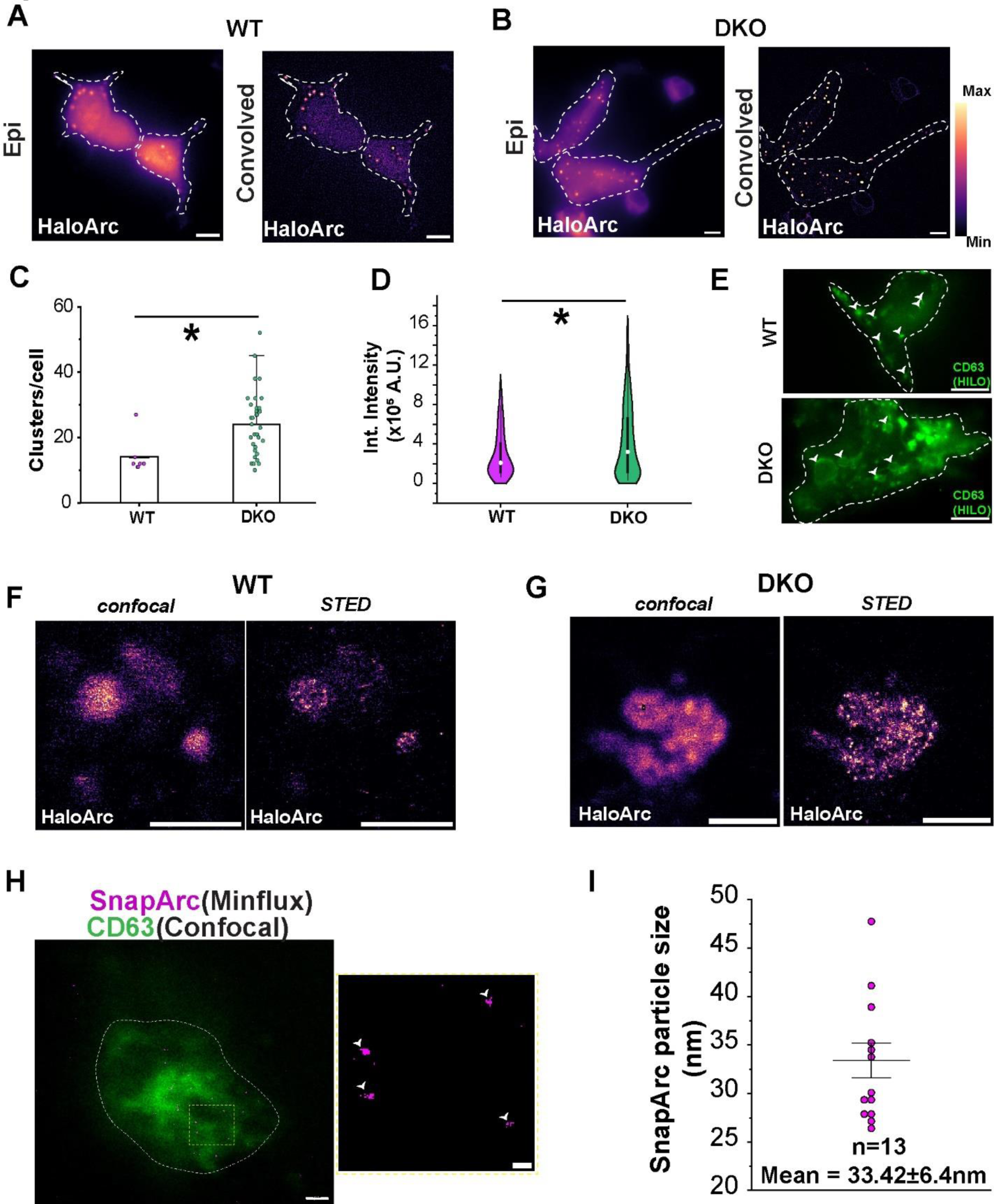
Erroneous MVB machinery alters the HaloArc phenotype in cells. A) Representative epifluorescence and convolved image of WT HEK293T cells expressing HaloArc. B) Representative epifluorescence and convolved image of DKO HEK293T cells expressing HaloArc. Scale bar in A and B: 2 µm. C) Quantification of the number of halo-Arc clusters in WT (pink) and DKO (green) HEK293T cells. D) Quantification of integrated fluorescence intensity of HaloArc clusters in WT (pink) and DKO (green) HEK-293T cells. E) A representative HILO image of WT HEK293T cell (top) and DKO HEK293T cell (bottom) expressing CD63-sfGFP, an MVB marker showing localization to intracellular vesicles. Scale bar in E and F: 5 µm. F) Representative image showing HaloArc clusters in a WT-HEK293T cell in confocal and STED mode showing the inner puncta of a single HaloArc cluster. G) Representative image showing an enlarged HaloArc cluster in a DKO-HEK293T cell in confocal and STED mode showing a higher number of inner puncta of a single HaloArc cluster. Scale bar in F) and G): 2 µm. H) Left, a cellular vesicle showing the expression of sfGFP-CD63 (confocal, green) and SnapArc (MINFLUX, magenta) labeled with Alexa-647 Snap ligand for MINFLUX imaging. Scale bar in H (left): 500 nm. Right, enlarged ROI to show multiple SnapArc capsids localized within CD63+ vesicles, an MVB. Scale bar in H (right): 50 nm. I) Size quantification of Arc particles captured using Minflux. (Mann-Whitney test; *: p<0.05).

Because of the limited spatial resolution of the epi-illumination fluorescence microscope, we were not able to resolve individual intraluminal vesicles within MVB. To address this issue, we applied STED super-resolution imaging to resolve fluorescently labeled Arc in fixed HEK-293T cells expressing HaloArc. Arc was localized by direct halo staining with JF-646 HaloTag ligand, an optimized fluorophore for STED microscopy. Arc puncta were first identified in the confocal mode, where no individual intraluminal vesicles could be teased apart. However, in the STED mode, we resolved multiple smaller puncta, likely individual intraluminal vesicles, within each MVB. Such a pattern was observed in both WT and DKO cells (**Fig. 6F-G)**. These results suggest that the RalA/RalB DKO primarily affects the membrane fusion of MVB with the plasma membrane but has minimal effect on the entry and assembly of Arc capsids into an MVB.

In order to resolve individual Arc capsids inside MVBs, we applied MINFLUX superresolution imaging to resolve fluorescently labeled Arc in fixed HEK293T cells expressing SnapTag-Arc and CD63-sfGFP. SnapTag-Arc was resolved in the MINFLUX mode by labeling SnapTag with Alexa-647, whereas CD63-sfGFP was resolved in the confocal imaging mode to label MVB. As expected, MINFLUX resolves that SnapTag-Arc forms multiple smaller puncta, likely individual intraluminal vesicles, within each MVB (**Fig. 6H**). The average size of Arc clusters resolved by MINFLUX was 33.4 ± 6.4 nm (n=13 particles, **Fig. 6I**), which matches the size of the reconstituted Arc capsids measured by TEM (**Fig. 3D-E**). These results suggest that Arc capsids could be encapsulated with intraluminal vesicles of MVB.

### RalA and RalB double knockout impairs Arc-mediated intercellular RNA transfer

To determine if defective MVB biogenesis leads to impaired Arc capsid release, we used Spectradyne’s nCS1 Particle Analyzer to measure the concentration and size of EV harvested from the conditioned medium of wild-type and RalA/RalB DKO cells, which were transfected with the same amount of HaloArc. nCS1 measures both endogenous (Arc-free) and Arc-containing EVs extracted from the conditioned medium in HEK293T cells. Compared to non-transfected HEK293T cells, HaloArc expression in WT cells boosted EV release by 2-fold, indicating Arc expression enhances EV secretion. In DKO cells, the basal level (Arc-free) EV concentration is significantly reduced, indicating a reduction of endogenous EV reduction. Although HaloArc expression still enhanced EV concentration in DKO cells, the absolute EV concentration is reduced by more than 50% compared to WT cells (**Fig. 7A**). HaloArc EVs in WT cells are larger than endogenous EVs, adding populations in the range of 100-250 nm in diameter. The larger EV size agreed with our STED images of the extracellular vesicles (**Fig. 1D, S1D**). In DKO cells, this portion of larger EVs disappears (**Fig 7B**).

**Figure 7.**
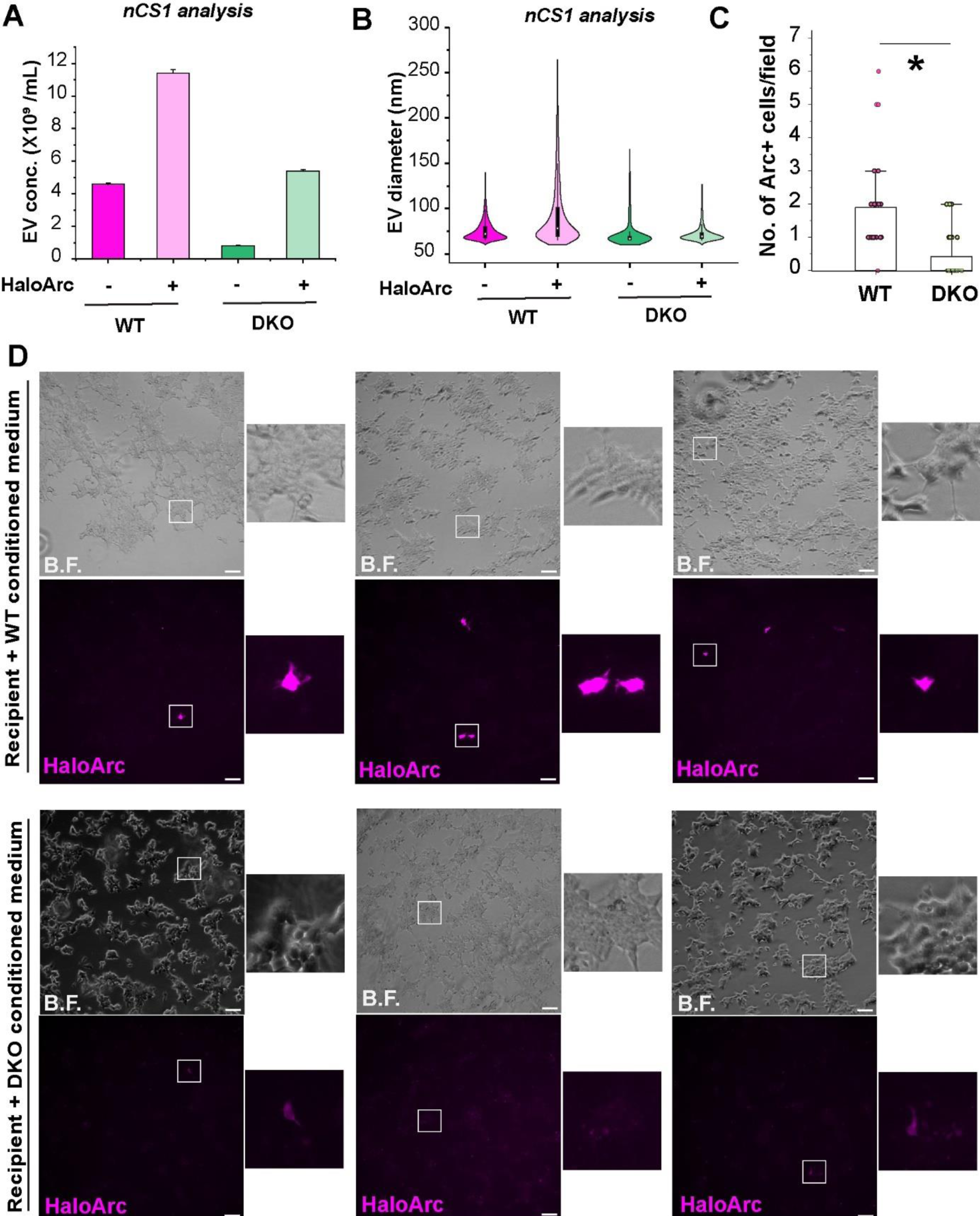
HaloArc exocytosis between cells depends on the MVB machinery. A) Comparison of EV concentration of WT and DKO cells with and without HaloArc expression measured on nCS1. B) Comparison of the EV size distribution of WT and DKO cells with and without HaloArc transfection measured on nCS1 showing DKO EV size does not change upon HaloArc expression. C) Transfer quantification of the donor-recipient assay performed using WT and DKO cells expressing HaloArc, showing transfer in recipient cells from DKO (n= 27 images) media is significantly lower than from WT (n= 29 images) media (Mann-WhitneyTest; *: p<0.05). D) Representative epifluorescence images (enlarged ROIs to the right) of recipient cells treated with HaloArc expressing donor cell media from WT (top) and DKO (bottom) cells, showing minimal transfer from DKO cell media. Scale bar: 100 µm.

We further confirmed that Arc-mediated intercellular RNA transfer is defective in the DKO cells using the donor-recipient RNA transfer assay. An identical amount of conditioned medium from WT and DKO donor cells expressing HaloArc was applied to two wells of equally plated HEK293T recipient cells. The transfection efficiency between WT and DKO donor cells is comparable. However, Arc expression in the recipient cells was significantly lower when incubated with the conditioned medium from DKO cells compared to that from wild-type cells, indicating defective intercellular Arc RNA transfer and protein expression in the recipient cells (**Fig. 7C-D**). Cells positive for Arc fluorescence from DKO cell media had a significantly lower fluorescence intensity compared to WT cell media (**Fig. S6A**).

## Discussion

In this work, we combined biochemical analysis, genetics, cell biological assay, live-cell, and superresolution imaging to define the intracellular axis of Arc capsid assembly and trafficking. Results suggest that the assembly and trafficking of Arc capsids is mediated through the endosomal-MVB pathway. Intriguingly, Arc capsid assembly and endosomal entry can be regulated by PI3P. Purified Arc proteins are associated preferably with PI3P. In mammalian cells, Arc colocalizes with FYVE, a PI3P marker. The high affinity between Arc and PI3P could recruit Arc proteins to endosomes or help with Arc oligomerization. Prolonged expression of FYVE reduces the average number of Arc puncta in cells, likely because of the competitive binding between FYVE and Arc to PI3P. Within the early endosomes, the protein undergoes sorting processes, likely involving specific protein-protein interactions or recognition motifs. This sorting directs the protein-containing vesicles toward late endosomes. Upon endosomal maturation, Arc capsids are packaged into individual intraluminal vesicles within MVB. As the MVBs mature, they migrate towards the cell periphery. Fusion between MVB and the plasma membrane releases Arc capsids as extracellular vesicles (**Fig. 8**).

**Figure 8.**
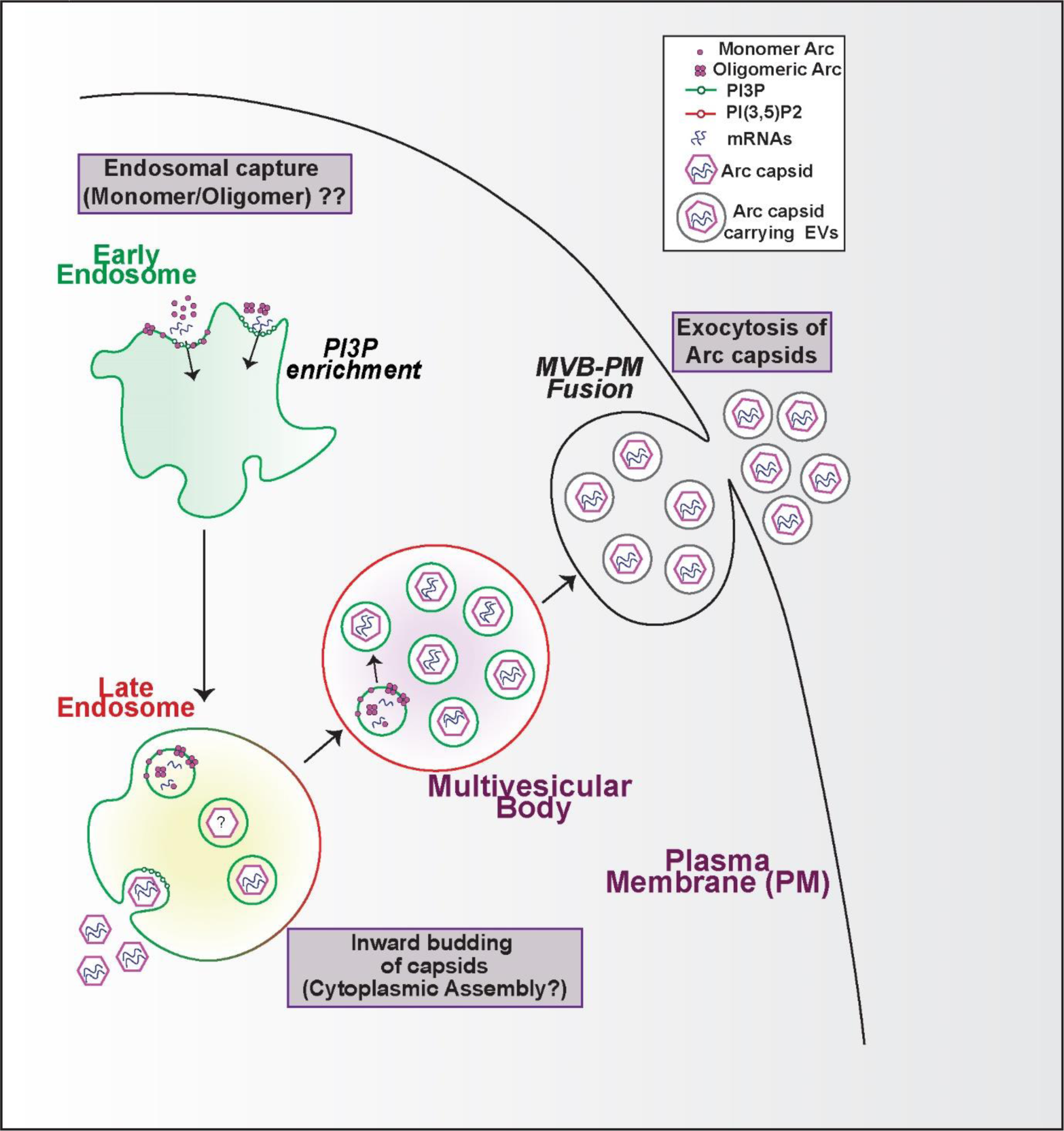
PI3P mediates Arc capaids trafficking and secretion through the endosome-MVBs pathway. Arc protein is targeted to the limiting membrane of the early endosome through its interaction with PI3P (green) and sorted through the axis of the late endosome and MVB. Endosomal PI3P could facilitate Arc oligomerization. Arc capsid assembly could occur either inside the cytoplasm or, more likely, within the intraluminal vesicles of MVB. Upon membrane fusion between MVB and the plasma membrane, extracellular vesicles containing Arc are secreted.

It is unclear if Arc enters endosomes as a complete capsid or incomplete assembly. We speculate that full capsid assembly is more likely to occur after endosomal entry due to the enrichment of Arc proteins. Live-cell imaging shows Arc puncta intensity is lower when colocalized with Rab5-positive early endosomes compared to Rab7- and CD63-positive MVBs. ILV entry of Arc may involve the Endosomal Sorting Complex Required for Traffic (ESCRT) proteins (42). Conventionally, ESCRT components 0, I and II help in the clustering of cargoes, curving the membrane and inducing their entry into the Intraluminal vesicles inside the MVBs. An ESCRT-independent ILV entry also exists that involves ESCRT-III, ALIX, and the Syntenin complex (43).

Trafficking through the endosomal-MVB pathway implies Arc capsids utilize distinct sorting signals to differentiate from other ILVs, which typically undergo degradation when MVB fuses with the lysosome. We speculate that the Rab family proteins, such as Rab11, that control endosome-to-plasma membrane recycling, could mediate Arc sorting. It has been suggested that Rab11 could promote cargo loading into the EVs or protect cargoes from lysosomal degradation. Recycling endosomes can populate neuronal EV precursor populations (44). Specifically, the loss of Rab11 reduces Arc levels in *drosophila* EVs (15).

PI3P is enriched in the limiting membrane of early endosomes. The high affinity of Arc to PI3P could facilitate the initial enrichment and recruitment of Arc to the early endosomal membrane. It is currently unclear which domain of Arc protein interacts with PI3P and if the binding to endosomal membrane is post-translationally regulated or constitutive. Stahelin L.V. and co-workers previously demonstrated that the PI3P interacting domain, FYVE, can bend the endosomal membrane, thereby increasing its residence time with the endosomes (45). It is conceivable that Arc binding also induces local changes in the endosomal membrane, which allows the entry of oligomeric Arc into the lumen of the endosomes.

PI3P could also play a role in mediating Arc capsid assembly. Biochemical analysis revealed that mammalian Arc consists of a positively charged N-terminal (NT) domain and a negatively charged C-terminal (CT) domain, separated by a flexible linker (46). In vitro experiments showed that Arc capsid assembly is mediated through the N-terminal domain, particularly the second helical structure, Coil-2 (47). Capsid assembly may start with Arc oligomerization on PI3P-enriched early endosomes. Such Arc oligomerization in the reducing cytosolic environment is possible because previous work showed that the five cysteine residues (Cys34, Cys94, Cys96, Cys98, and Cys159) did not affect the Arc-Arc interaction, suggesting that oligomerization does not require disulfide bond formation (47). Compared to monomeric Arc, Arc oligomers bind to PI3P with higher avidity, which could gate the initiation of precise Arc capsid assembly by ensuring sufficient capsid building blocks. Oligomeric Arc could be invaginated from the limiting membrane of MVB and become intraluminal vesicles, where full capsid can be assembled as the endosomes mature.

The assembly and trafficking of capsids of Arc and HIV Gag protein present intriguing parallels and distinctions, offering insights into commonalities and unique mechanisms underlying these processes within biological systems. Similar to other viral Gag proteins, Arc proteins self-assemble into capsids in the presence of phosphate or nucleic acids. Both Arc and HIV rely on multimerization for capsid assembly. Arc, a neuronal protein, forms virus-like capsids through self-oligomerization, a process akin to the assembly of retroviral capsids. This self-assembly ability mirrors the structural arrangement observed in the assembly of HIV capsids, where the Gag polyprotein undergoes multimerization to form the viral capsid. Both Arc and HIV capsid shells serve as a protective convoy vehicle that protects the encapsulated genetic materials.

Phospholipids seem to play a crucial role in HIV and Arc capsids assembly in mammalian cells. It is known that HIV capsid proteins reconstitute capsid-like structures in vitro but form only 25-30 nm in size unless mixed with inositol phosphates like IP5 and PIP3, which restores the 100-120 nm HIV capsids comparable to their sizes assembled in cells (48). In addition, the HIV gag specifically restricts PI(4,5)P2 and cholesterol on the plasma membrane for its assembly (49). Indeed, Linwasswe O. W. and coauthors showed that the HIV-gag assembly site is regulated by the subcellular distribution of cholesterol (50). In this work, we found that phospholipids regulate Arc capsid assembly in mammalian cells. However, Arc has a higher affinity to PI3P, an endosome-specific phospholipid.

In contrast to the asymmetric cone formed by HIV capsids, Arc forms a spherical capsid with a diameter of 30 nm (**Fig. 3D and 6H-I**), consistent with previous findings (14). This difference might result from distinct monomer interface and oligomeric units formed by HIV capsid protein and Arc. HIV can form slightly different pentameric and hexameric assemblies, both serving as building blocks that tile capsid surfaces. Another difference is that HIV capsids are commonly believed to be released at the plasma membrane, whereas we found that Arc capsids secrete through the endosomal-MVB signaling axis. However, HIV capsid release might be cell-type specific. Ono A. et al. previously compared HIV assembly of MA mutants in different cell types and discovered that HIV can also utilize the MVB pathway for its assembly and egress upon removal of plasma membrane targeting basic MA domain residues. In fact, monocyte-derived macrophages targeted the WT gag to the intracellular vesicles, which was confirmed by colocalization with CD63 and Rab7 (51). The MVB pathway utilization has also been the murine leukemia virus (MLV) replication (52).

Synaptic development is an important function of the neuronal IEGs and corresponds to their link with various neurological conditions when dysregulated. Arc is different from other IEGs in that Arc is not a transcription factor but rather a major signaling hub at distinct subcellular compartments such as synapses (53–55), axons (56), or the nucleus for transcriptional regulation (57) and chromatin modification (58). The recently discovered Arc function in regulating intercellular RNA transfer requires a stable infrastructure where Arc can go through intracellular assembly into a capsid and intercellular delivery of genetic materials. On the one hand, genetic materials encapsulated in the capsids sample the intracellular state at the moment of release. On the other hand, these materials could define the Arc’s remote functionality in the recipient neurons yet to be discovered.

The involvement of PIP lipids in Arc neuronal intercellular communication is intriguing but makes sense, considering PIP lipids have a major role in neuronal function, particularly in modulating synaptic strength. PI3P is an important signaling lipid and is converted to its important metabolite PI(3,5)P2 using PIKfyve (38, 59). PI(3,5)P2 levels bidirectionally control the synaptic strength, with high PI(3,5)P2 causing synaptic pruning and low PI(3,5)P2 levels causing synapse strengthening. PI(3,5)P2 repression also reduces AMPAR endocytosis, a process also directly controlled by Arc protein (60). Interestingly, the PI3P-producing enzyme Vps34 is highly expressed in the central nervous system neurons and is highly localized at the dendritic spines. Loss of Vps34 is associated with loss of dendritic spines and gliosis and neurodegeneration in aged mice (61), supporting the idea that higher PI3P levels might be required during synaptic strengthening, such as long-term potentiation (LTP). The neuronal function might depend on the PI3P: PI(3,5)P2 ratio as several genes related to the synthesis of PI(3,5)P2 from PI3P have been implicated in critical neurological conditions like epilepsy, neuropathy and neurodegeneration. As an early endosome and MVB marker, PI3P regulates the recruitment of PH-, PX- and FYVE domain-containing proteins to MVB compartments (62). Initial membrane attachment of proteins can induce dimerization and further penetration of hydrophobic amino acids into the intracellular membrane (45, 62).

We noted that Arc puncta populated across the dendrites in cortical neurons (**Fig. 2H**). In supporting this idea, electron microscopy also resolved immunogold-labeled Arc in the dendritic spines (63). As shown in this work, the release of Arc capsids can go through the MVB pathway, mediated by Ral GTPases. Indeed, neuronal MVBs have been implicated in various aspects of neuronal system regulation via cellular material exchange (64, 65). Electron microscopy has revealed that the dendritic shafts of the neurons are enriched with MVBs, with a fraction of spines of the glutamatergic synapses also carrying some MVBs (66, 67). Strikingly, somato-dendritic compartments carry about 50 times more MVBs than axons (68). MVBs migrate toward synapses in response to synaptic activity and learning behavior (e.g., water maze) in rats (69). Chronic stress, on the other hand, reduces the association of MVBs with PSD (69). The dendritic membrane undergoing membrane fusion with MVB could be a potential site for Arc capsid release. However, a direct visualization of such an event is yet to be elucidated.

Interestingly, a recent study has shown another synaptic protein, gephyrin, found at the inhibitory synapses, also clusters in response to PI3P in cultured hippocampal neurons, thereby determining the strength of inhibitory GABAergic synapses (39). In line with this idea, defective PI3P synthesis could lead to improper Arc release and over-accumulation of Arc at the dendrites, causing silencing and even loss of the spines. This loss remains permanent and could lead to progressive degeneration in the brain (61). Evidently, AD patients have a PI3P deficiency, and PI3P levels are genetically linked with ALS and PD (70). Understanding the cellular machinery orchestrating Arc capsid assembly and release promises to shed light on the commonality and distinction of trafficking pathways between structurally resembled capsid proteins, as well as intercellular communication between distant neurons.

## Materials and Methods

### Plasmid construction

The HaloArc sequence was cloned from pRK5-mHalo-Arc plasmid, a gift from Prof. Jason Shephard, University of Utah. The HA sequence was inserted into the multiple cloning site (MCS) of an empty pCS2+ vector (NovoPro) using InFusion cloning after double digestion with FD-EcoRI (Thermo) and FD-NotI (Thermo). HaloTag was replaced with sfGFP using InFusion (Takara) assembly by cutting out the HaloTag using FD-PacI (Thermo) and FD-BcuI (Thermo). HaloTag and sfGFP were independently amplified from the HaloArc and sfGFP-Arc plasmids, respectively and cloned into an empty pCS2+ vector using InFusion assembly. All other plasmids used in this paper were either purchased and used directly or cloned out from the Addgene plasmid and used with specific. All plasmid information has been provided in **Table S1**. All information regarding the reagents has been provided in **Table S3**.

### Lentivirus production

The HA sequence was inserted into a pLJM1 backbone after double digestion with FD-NheI (Thermo.) and FD-EcoRI (Thermo.) using InFusion cloning (Takara). For neuron knockdown, two shRNA encoding sequences (https://sirna.wi.mit.edu/) were inserted into FD-EcoRI and FD-BshTI digested pLKO.1 vector using a similar strategy as for KO plasmids (see above). Low-passage (< p10) HEK293T cells were plated in growth media without penicillin or streptomycin and incubated overnight. A transfection solution of 3:2:1 of the plasmid of interest, psPAX2 (packaging), and pMD2.G (envelope) plasmids diluted in Opti-MEM was created, along with a 3:1 dilution of PEImax diluted in Opti-MEM. After 5 minutes, the two solutions were combined, incubated for 30 minutes at room temperature, and added dropwise to cells. After exactly 16 hours post-transfection, the media was changed to complete growth media. After 24 hours of incubation, the media was harvested and stored at 4 °C and replaced with fresh growth media. After another 24 hours of incubation, the media was harvested again and combined with the previously stored media. The media was centrifuged for 5 minutes at 500 g and filtered through a 0.45 μm syringe filter (Millipore). The media was concentrated with a UV-sterilized Amicon ultra-15 100 kDa centrifugal filter until the solution was concentrated by 30 times its original volume. Aliquots of this concentrated media were made and stored at -80 °C. All information regarding the reagents has been provided in **Table S3**.

### Protein purification and characterization

Arc protein was overexpressed using the pGEX-6p1-rArc construct, a gift from Prof. Jason D. Shepherd, University of Utah. Briefly, house-made chemically competent BL21(DE3) cells were transformed with the plasmid. A starter bacterial culture was grown overnight at 37 °C in Luria broth LB (Fisher) supplemented with ampicillin (Fisher). Starter cultures were used to inoculate large-scale 500 mL cultures of LB. Once the culture was grown to OD600 of 0.6-0.8 at 37 °C at 150 rpm, the culture was moved to a 19 °C incubator set at 150 rpm for 16-20 h. Cells were pelleted at 5000 g for 15 min at 4 °C and cell pellets were resuspended in 30 mL lysis buffer (500 mM NaCl, 50 mM Tris, 5% glycerol, 1 mM Dithiothreitol DTT, 1 mM Phenylmethylsulfonyl fluoride PMSF, pH 8.0 at room temperature (RT)). Resuspended cells were flash-frozen in liquid nitrogen and stored at -80 °C overnight. The next day, frozen pellets were thawed quickly at 37 °C and brought to a final volume of 1 g pellet:10 mL lysis buffer. Lysates were then sonicated for 5 minutes with 2.5 sec on, 2.5 sec off at 40% amplitude and pelleted for 45 min at 21,000 g. Cleared supernatants were then passed through a 0.45 µm filter and incubated with pre-equilibrated Glutathione Sepharose 4B affinity resin (Cytiva) in a gravity flow column overnight at 4 °C on a rolling platform. Bound protein was then washed twice with two column volumes of lysis buffer, re-equilibrated with cleavage buffer (150 mM NaCl, 50 mM Tris, 1 mM Ethylenediamine tetra-acetic acid EDTA, 1 mM DTT, pH 7.2 at RT), and cleaved on-resin overnight at 4 °C with 100uL PreScission Protease (GE). Elution was collected the next day with 5mL elution buffer (50mM Tris, 10mM reduced L-glutathione, pH 8.0). Cleaved proteins were run on an S200 size exclusion column (Cytiva) to separate the cleaved protein, and peak fractions were pooled. Protein was concentrated and buffer exchanged in TBS buffer (50 mM Tris, 150 mM NaCl, 1 mM DTT, pH 7.4). Average yields for purified proteins were ∼10 mg per 500 mL of cell culture. All information regarding the reagents has been provided in **Table S3**.

### PIP-Strip lipid overlay assay

Purified Arc protein was spun at 13000 g for 10 minutes at 4⁰C prior to the PIP strip assay. A 0.5uL protein was spotted and dried out completely in the corner of the PIP strip (Echelon Bio.) to serve as the positive control. Later, the strip was incubated in a blocking solution (5% milk in TBS, pH 7.2) on a rocking platform for 1 hour at RT. Protein was diluted in fresh blocking buffer (1ug/mL). Diluted protein was added to the strip and incubated for 2 hours at RT on a rocking platform. The strip was washed thrice with TBS. Primary Arc antibody (Synaptic Systems) was diluted in blocking solution (1:1000). The strip was incubated with Arc antibody for 1 hour at RT on a rocking platform. The strip was washed again three times with TBS and incubated with HRP-conjugated secondary antibody (1:10000) for 1 hour at RT on a rocking platform. The strip was washed thrice and then imaged for chemiluminescence using freshly prepared Clarity ECL western substrate (Biorad). All information regarding the reagents has been provided in **Table S3**.

### HEK293T cell culture and transfection

HEK293T cells were cultured in Dulbecco’s Modified Eagle Media (DMEM, Corning) supplemented with 100 units/mL Penicillin, 0.1 mg/mL Streptomycin, 2 mM L-Glutamine and 10% fetal bovine serum (Sigma Aldrich). Cells were grown in an incubator maintained at 37°C and 5% CO2. Before transfection, cells were plated and allowed to stabilize overnight. Exactly 1 hour before transfection, cell media was changed to starvation media containing 1% serum. The transfection mix was prepared by diluting plasmid DNA in fresh DMEM without supplements. PEImax (PolyScience) was used to carry out transfection and was added as per the vendor’s instructions. Media was changed back to growth media 3-5 hours post-transfection. About 24-48 hours after media change, cells were analyzed for specific downstream applications. All information regarding the reagents has been provided in **Table S3**.

### CRISPR knockout and monoclonal selection

Gene knockout was performed for RalA and RalB in HEK293T cells using the CRISPR-Cas9 technology. A total of 4 gRNAs were designed for both hRalA and hRalB genes using CHOPCHOP (https://chopchop.cbu.uib.no/). Briefly, each primer pair was annealed and consequently phosphorylated with T4-polynucleotide kinase (T4-PNK; NEB). Restriction digestion of the pLentiCRISPRv2 vector was performed using FD-Esp3I (Thermo) and dephosphorylation was done later using the Shrimp alkaline phosphatase (rSAP, NEB). An overnight ligation was performed by mixing digested vector and annealed gRNA inserts using T4 DNA ligase (NEB) at 4⁰C. The plasmid was then transformed into competent *E. coli* cells. All KO plasmids were directly transfected into the cells using PEImax. Exactly 16 hours after transfection, cells were recovered in growth media supplemented with 1.2 μg/mL puromycin. Cells were cultured in selection media for another 2 weeks. Monoclonal selection was performed with selected cells in a 10-cm dish. Colonies started to appear in about 10 days and were picked and transferred to individual wells of a 24-well plate. Double knockout was confirmed from reverse transcribed cellular RNA against hRalA and hRalB specific primers. All information regarding the reagents has been provided in **Table S3**.

### Primary Neuronal Culture and Lentiviral transduction

Coverslips were coated with 100 µg/mL Poly-D-Lysine (PDL) diluted in Borate buffer. The next day, PDL on the coverslips was aspirated out and washed twice in autoclaved Mili-Q water. Coverslips were left to dry inside the laminar airflow under UV for about 20 minutes until they were ready for plating. Brains from E18-E19 rat embryos were placed in a large Petri dish containing 1× slice dissection solution (SLDS) at 4°C. Cortices were dissected out of the brains, and about 2 mL protease enzyme solution was added into the cortices and incubated inside an incubator at 37 °C for 30 minutes for dissociation. The enzyme solution is removed gently using a pipet, and a 3 mL plating medium is added into a conical tube containing the cortices. Tissues are dissociated into single cells using polished glass pipets. Cells are then counted on a hemocytometer, and about 500,000 neurons are plated per coverslip in a plating medium. Media was changed to neurobasal (NB) media after 3 hours of plating. Half of the NB media was changed every 3 days to avoid loss of growth factors.

Neuron transduction was performed between DIV6 and 7. Briefly, half of the NB media was removed, and an appropriate content of the virus was added to the neurons. After 3 hours of incubation, the NB media containing the virus was removed and stored NB was added back to the transduced neurons with equal volume of fresh NB. Neurons were maintained normally for 6-12 days to allow expression. Transduced cells to be imaged were labeled with 200 nM of JF-549 HaloTag ligand and were kept in ACSF during the destaining step. Labeled neurons were fixed with 4% paraformaldehyde, immunolabeled for MAP2 and mounted onto the glass slides using the ProLong Gold antifade reagent with DAPI (Thermo.). All information regarding the reagents has been provided in **Table S3**.

### Fluorescence labeling of HaloArc and Immunofluorescence

Transfected cells were stained with JF-549 tagged HaloTag ligand (Promega) and Vybrant DiO staining solution (Invitrogen) in phenol red-free DMEM for about 15-20 minutes at 37⁰C at a final concentration of 200 nM and 1 µM respectively. For HEK293T labeling, cells were washed twice with Dulbecco’s phosphate-buffered saline with Calcium and Magnesium (Corning) to minimize cell detachment. For primary neuron labeling, neurons cultured on coverslips were stained in the NB media. Neurons were washed and destained in artificial cerebrospinal fluid (aCSF) media with or without high KCl (50 mM) for 1 hour. Neurons were then fixed in 4% paraformaldehyde and 3% sucrose in PBS for 15 minutes on ice. After fixation, neurons were permeabilized in 0.2% TritonX-100 in PBS for 10 minutes. Neurons were blocked in 1% BSA in PBS for 30 minutes at RT. Blocked neurons were incubated with primary MAP2 antibody diluted 1:2000 in blocking buffer for 16 hours at 4 °C. After washing with PBS, cells were incubated with 1:1000 dilution of Alexa-647 labeled secondary antibody for 2 hours at RT. Coverslips were mounted onto glass slides using the Prolong Gold Antifade mounting medium with DAPI (Thermo.) and cured overnight in the dark. All information regarding the reagents has been provided in **Table S3**.

### Epi-illumination Microscopy

Cells were finally imaged in serum and phenol red-free DMEM (Gibco) at 40× air and 100× oil objective using an epi-illumination inverted fluorescence microscope (Leica DMI8) equipped with both the objectives and a light-emitting diode illuminator (SOLA SE II 365). JF-549 fluorescence was detected using the mCherry filter cube (Leica, excitation filter 560/40, dichroic mirror 585, and emission filter 592/668) with an exposure time of 500 ms for JF549 (red), 200 ms for Alexa 647(far red), 200 ms for DAPI (blue) and 50 ms for DiO (green), and 2×2 binning.

### HILO fluorescence imaging

Transfected cells cultured on coverslips were stained with JF-549 HaloTag ligand and were imaged under a custom-built HILO microscope. Cells were tracked for 20 s using an integration time of 100 ms in HILO mode. A continuous wavelength (561 nm, Spectral physics) laser was used as the light source. An inverted microscope (IX73, Olympus) equipped with a 100× oil immersion TIRF objective (Olympus, PlanApo, 100×, NA 1.49, oil immersion) was used. The laser beam was then expanded, collimated to about 35 mm, and directed into the microscope by a lens (400 mm focal length, Thorlabs LA1725A). The incident light was directed through the objective via an exciter (FF01-482/563-25) and a dual-band dichroic filter (Di01-R488/561−25 × 36). The cellular fluorescence was collected by the same objective, passing an emitter (FF01-523/610-25) and captured by an electron multiplying charge-coupled device (EMCCD) camera (iXon U797, Andor Technology). Arc clusters were detected as bright intracellular clusters. HaloTag transfected cells, serving as a negative control, were labeled and tracked with an integration time of 10 ms achieved by reducing the size of the field to 256×256 pixels.

For two-color imaging in the HILO mode, the incoming fluorescence from the microscope was split into red and green light using the Opto-Split II (Cairn Research) setup installed between the camera and the light path (**Fig. S3A**). Fluorescence was separated by two filter sets (FF03-482/563-25 for green emission and FF02-523/610-25 for red emission. Emitted light was split using a 575LP (Di R488/561). Co-transfected cells were prepared and illuminated concurrently with 488 and 561 nm lasers. The split images were visualized on a split screen on a computer. Tracking was performed with an integration time of 100 ms for 20 s. Images were later cropped and merged in FIJI (https://fiji.sc/). All videos were analyzed for colocalization, intensity and diffusion coefficient calculations (see below).

### Super-resolution imaging (STEDYCON and MINFLUX)

For STED imaging, cells and EV samples were labeled with 200 nM JF646 HaloTag ligand. EVs were co-stained with 1 µM DiO to locate them. The samples were fixed and mounted onto glass slides using a DAPI-free mount. Samples were initially imaged in confocal mode, and then a selected ROI was run on the STED mode using the Abberior STEDYCON setup.

For MINFLUX imaging, cells were co-transfected with SnapArc and CD63-sfGFP constructs as previously. Briefly, 24 hours post-transfection, cells were fixed for 30 s in 2.4% paraformaldehyde. Cells were immediately permeabilized with 0.4% TritonX-100 for 3 min and fixed again in 2.4% paraformaldehyde for 30 min. Cells were quenched with 100 mM NH4Cl prepared in PBS. After washing twice with PBS, cells were blocked with Image-IT signal enhancer (Thermo.) for another 30 min. Alexa-647 Snap dye (NEB) was diluted in the staining solution (1 mM DTT+0.5% BSA in PBS) to a final concentration of 100 mM and added to cells for 50 min. Cells were washed 2-3 times in PBS. Shortly before imaging, coverslips were treated with gold beads (BBI solutions) for 10 min. Coverslips were washed multiple times and were inverted onto a cavity slide containing imaging buffer (50 mM Tris-HCl, 10 mM NaCl, 10% glucose, 64 μg/mL catalase, 0.4 mg/mL glucose-oxidase,10 mM cysteamine, pH=8.0) and sealed with a two-component silicone glue (Picodent, Twinsil). Cells were first imaged using the 488 nm and 640 nm lasers in the confocal mode to find a region of interest. The region of interest was then cropped from the whole image and captured in a new window with a pixel size of 20 nm. This selected ROI was used to carry out MINFLUX imaging using the 640 MINFLUX laser. Data was collected for about 30 min, and a 405 nm laser was used to re-activate the fluorophores. Images in confocal (for CD63) and MINFLUX-mode (SnapArc) were overlayed after resizing in FIJI.

### Donor-recipient transfer assay

HaloArc transfer was assayed by transferring conditioned media from HaloArc-transfected or untransfected cells (donor) to untransfected cells (recipient). Briefly, donor cells were cultured on a 10-cm dish and transfected with 10 μg HaloArc plasmid. Cells were washed twice with DPBS to remove any excess plasmid-PEI aggregates after 5 hours of transfection, and fresh growth media was added. Conditioned cell media from transfected donor cells was harvested 24 hours later and spun at 500 g for 5 minutes to remove any cell debris. Cleared supernatant was then treated to the recipient cells cultured on 12-well plates and incubated for another 24 hours. Transfected donor cells were concurrently stained with JF-549 HaloTag ligand and imaged under epi-illumination to confirm expression. Recipient cells were stained after donor media treatment across the entire well. Cells showing HaloArc expression were counted per field and analyzed to quantify intercellular RNA transfer. All information regarding the reagents has been provided in **Table S3**.

### Western Blot

Transfected cells were lysed 24 hours post-transfection using cold lysis buffer (1× RIPA, 1× protease inhibitor cocktail). Cell lysates were centrifuged at 17,000 g for 10 minutes at 4 °C. Protein in the supernatant was quantified on a plate reader using the Bradford assay (Sigma Aldrich). All samples were prepared in loading dye (NU-PAGE dye + 5mM BME) and boiled at 99 °C for 5 minutes. Equal amounts of denatured lysates were loaded on a 12% SDS gel and ran with SDS running buffer (Biorad) for 90 minutes at 100 V. An overnight transfer was performed to capture proteins onto PVDF membrane (Biorad) with transfer buffer (25 mM Tris-base, 192 mM Glycine, 10% Methanol). The membrane was blocked in a blocking buffer (5% milk in Tris-base, 1%Tween-20; called TBST) and was incubated with primary antibodies overnight at 4⁰C. After washing with TBST, the membrane was treated with HRP-conjugated secondary antibody and later stained and imaged for chemiluminescence using freshly prepared Clarity ECL substrate (Biorad). All information regarding the reagents has been provided in **Table S3**.

### Reverse transcription PCR

Total cellular RNA was extracted using the GeneJet RNA extraction kit (Thermo). A total of 1 μg of cDNA was prepared using the M-MulV reverse transcriptase (NEB) as per vendor instructions. A total of 10 ng cDNA was used to run qPCR using the SYBR green master mix (Applied Biosystems). All gene-specific primer sequences have been provided in **Table S2**. The reactions were run on an Applied Biosystems quantitative PCR machine, and all the acquired Ct values were analyzed. In all the samples, the *36B4* gene served as the internal control gene. To confirm the CRISPR gene knockout for RalA and RalB in the monoclonal cells, a regular PCR was run on a thermocycler using Taq polymerase master mix (Promega). Gene knockout was thereby confirmed by gel electrophoresis. All information regarding the reagents has been provided in **Table S3**.

### Extracellular vesicle isolation and downstream applications

HEK293T cells were cultured in standard growth media for 24 hours post-transfection before media harvest. EV-depleted growth media was prepared using a 100 kDa cut-off ultrafiltration filter (Amicon) by centrifugation of FBS at 3,000 g for 55 minutes before media preparation. For nCS1 experiments, transfected cells were cultured in EV-depleted growth media for an additional 24 hours before media harvest. The Capturem EV isolation mini kit (Takara) was used following the manufacturer’s instructions. Briefly, the conditioned media was spun at 500 g for 5 minutes and then at 20,000 g for 30 minutes. The collected supernatant was loaded onto the pre-clearing columns to remove larger particles larger than 800 nm. The cleared media was then concentrated using a 100 kDa cut-off ultrafiltration filter by spinning at 3000 g until the final volume was reduced to about ∼1mL. The concentrated media was then loaded onto the purification column, and EVs were eluted in the elution buffer.

The concentration and size of EV was measured using Spectradyne nCS1. Eluted EVs were buffer exchanged with freshly prepared, 0.22 μm-filtered 0.1%-PBST (pH=7.4). A 5 μL sample was applied to the Spectradyne C-400 cartridges (diameter range 65-400 nm). The samples were run on the nCS1 instrument until at least 5000 particles were analyzed. Peak data was filtered according to the guidelines in the Spectradyne nCS1 manual. False-positive particles are characterized as having transit times >100μs, diameters outside of 65 to 400 nm, a signal-to-noise ratio of <15, or a peak symmetry value outside of 0.2 to 4.0. Data and statistical analysis were performed using OriginPro (2021b).

Two-color fluorescence imaging of EV was performed by staining concentrated EVs with 5 μM DiO cell staining solution (Invitrogen) and 100 nM JF-549 HaloTag ligand (Promega) before the column binding step. They were fixed and imaged at 100× magnification or a 2D-STEDYCON setup at IGB of UIUC.

EV RNA was isolated from eluted EVs (with or without RNaseA treatment) using the TriZOL reagent (Invitrogen) as per the user manual. Isolated RNA was subjected to cDNA synthesis using random or poly-A primers. PCR was carried out using mArc cDNA specific primers (**Table S2**) using Taq DNA master mix. PCR samples were thereby run on an agarose gel to confirm the presence of HaloArc mRNA inside EVs.

Eluted EVs were lysed using RIPA buffer and incubated on ice for 5 minutes. Lysed samples were spun at 14,000 g for 15 min at 4 °C. The supernatant was later processed for western blot as before.

### Particle tracking and diffusion coefficient calculation

Particles were tracked from the HILO-acquired videos using the FIJI plugin TrackMate(71) (https://imagej.net/plugins/trackmate/). The *‘spots.csv’* file generated in TrackMate was used for diffusion coefficient analysis. Briefly, all the tracks were exported from the spots file to a custom-written Python script (**Software S1**), which calculated the particle diffusion coefficient using the Co-variance Estimation (CVE) method(72, 73). The diffusion coefficient (DCVE) was calculated as follows:

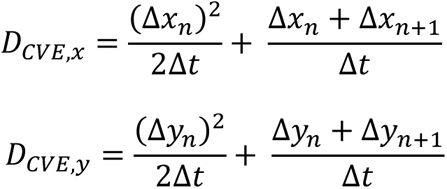

where Δxn is xn+1-xn in a trajectory time series where Δt is the integration time. The entire series of tracks was averaged out to get the final DCVE, as mentioned below.

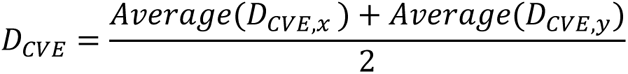

Particle fluorescence intensity was acquired as the maximum value of the intensity was acquired from each particle tracked by TrackMate. Co-localization for 2-color HILO videos was quantified by manual counting for all HaloArc clusters in the tracked videos.

### Negative-Stain Transmission Electron Microscopy of Arc capsids

Arc capsid assembly and negative staining were performed as done previously(14). Briefly, the purified Arc protein was incubated with either cellular RNA extracted from HEK293T cells or RNA (of Arc or GFP) that was transcribed *in-vitro* using MAXIscript SP6 Kit (Invitrogen) at 7.3% w/w RNA-to-Arc ratio (10 nucleotides of RNA to 1 molecule of Arc). Once the RNA was added, the sample was diluted in high phosphate buffer (500 mM NaPO4, 0.5 mM EDTA, 50 mM Tris, pH 7.5) to 0.25 mg/mL and incubated at room temperature for 2 hours. A Copper 200-mesh grid coated with Formvar and carbon (Electron Microscopy Sciences Cat# FCF200CU50) was used for negative staining. Grids were discharged for 25 seconds in a vacuum chamber at 15mA using the PELCO easiGlow 91000 Glow Discharge Cleaning System. A 5 µL sample was added to the grid. After 2 minutes, the excessive liquid was blotted using filter paper. The grid was dipped once in a 50-µL MiliQ water blob and then in a 50 µL 1% uranyl acetate blob on parafilm; the excessive solution was blotted gently in between with filter paper. Then, 5 µL of 1% uranyl acetate is added to the grid to stain for 30 seconds. The excessive stain was blotted and air-dried for 1 minute. All images were acquired under the JEOL JEM-1400 TEM in the Materials Research Laboratory Central Research Facilities, University of Illinois Urbana-Champaign. All information regarding the reagents has been provided in **Table S3**.

### Data Analysis, Statistics and Data Visualization

HaloTag and HaloArc intensity profiles were generated by plotting the fluorescence intensity across the length of the region of interest (ROI). Peaks in the intensity profiles correspond to the presence of Arc clusters. Colocalization was confirmed by plotting the intensity profiles of both channels into a single plot. HaloArc cluster parameters were calculated using the ‘Analyze Particles’ feature of FIJI. Images were initially convolved using the *‘Convolve’* filter in FIJI for background subtraction and feature enhancement. We then adjusted the threshold of the images using the ‘Threshold’ feature of FIJI until all the clusters were picked up as individual particles. Particles were then analyzed, and all the measured parameters were exported to OriginPro 2022 (https://www.originlab.com/2022) and plotted into histograms and violin plots. Datasets were compared using the Mann-Whitney U or the Kruskal-Wallis ANOVA non-parametric test as it does not assume the normal distribution of the datasets. A p-value of <0.05 was reported as significant and was shown using a ‘*’ between datasets.

## Author Contributions

K.M., H.Y., J.L., T.T.G. and K.Z. formal analysis, validation, investigation, and visualization; K.M., T.T.G and K.Z. methodology; K.M. and K.Z. conceptualization; K.Z. resources; K.M. and K.Z. Data curation; K.M. and K.Z. Software; K.Z. Supervision; K.Z. funding acquisition; K.M. and K.Z. writing-original draft; K.Z. project administration.

## Acknowledgments

We thank Dr. Jason D. Shephard, University of Utah for providing the HaloArc and GST-Arc constructs; Dr. Beth M. Stadtmueller and Sonya K. Bharathkar, UIUC, for SEC-AKTA usage; Dr. Milan Bagchi, UIUC for Spectradyne nCS1 usage; Material Research Laboratory Central Research Facility, UIUC for JOEL JEM-1400 TEM usage; Core facilities at the Carl R. Woese Institute of Genome Biology, UIUC for MINFLUX and STEDYCON usage; and Huaxun Fan, UIUC for help with primary cortical neuron culture. This work is supported by the National Institute of General Medical Sciences (R01GM132438) and National Institute of Mental Health (R01MH124827) of the National Institutes of Health, National Science Foundation (Award #: 2121003) and an STC-QCB center grant (K.Z.).

## Supporting Information

**Fig. S1:**
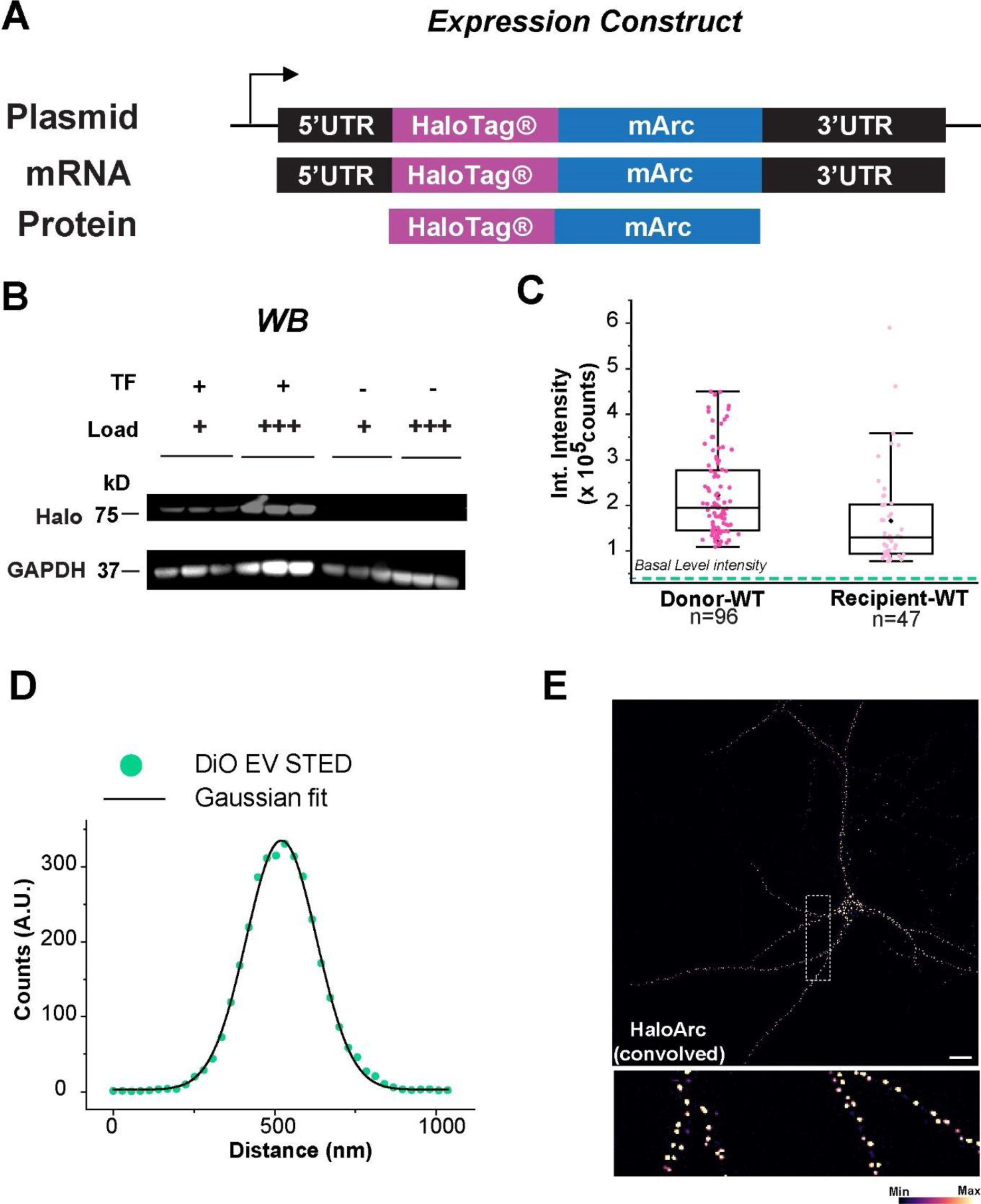
A) Schematic representation of the plasmid construct used for expressing mArc in HEK-293T cells. The plasmid carries an N terminal HaloTag and untranslated regions on both the 5’ and 3’ ends of the RNA. B) Western blot carried out using the cell lysate from transfected and untransfected cells. A 75 kDa band shows the expression of HaloArc in cells upon transfection. C) Box plot of fluorescence intensity quantification of cells in the donor-recipient assay. The green dashed line shows the basal intensity of untransfected cells. D) Intensity profile of and STED image of an EV (green dots) in Fig. 1D fitted with a one-dimensional Gaussian curve (black). E) A representative image of HaloArc transduced primary rat cortical neurons showing the HaloArc cluster intensity after convolution. Warmer colors indicate higher intensities. The marked ROI is enlarged (bottom) to show neuronal processes carry bright clusters of HaloArc. Scale bar: 20 µm.

**Fig. S2:**
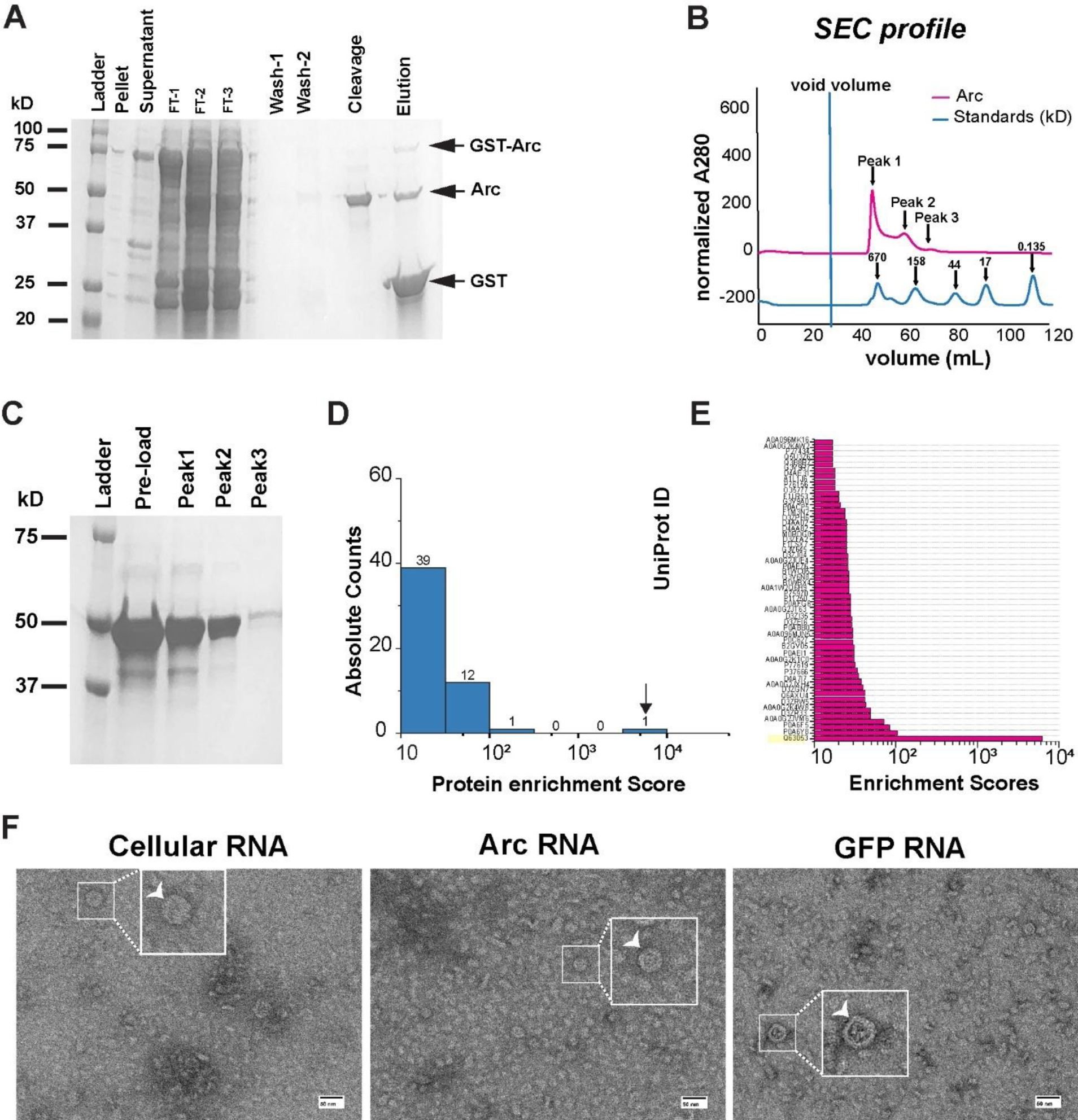
A) GST-Arc overexpression n bacteria and affinity chromatography protein gel showing GST-Arc (75 kDa) was present mostly in the supernatant fraction after lysis. GST cleavage results in the elution of pure Arc protein (45 kDa). B) SEC profile of pure Arc is run for size exclusion column overlapped with protein standards showing Arc elutes in 3 different sized oligomers (peak 1,2 and 3). C) Protein gel for different SEC fractions collected showing a single 45 kDa band on SDS-PAGE. D) Mass spectrometry results showing protein enrichment scores for the top 53 hits with p<0.05. E) Individual scores identified Arc (uniport ID: Q63053) as the top hit. F) TEM micrographs of capsids induced under high phosphate conditions with cellular RNA (left), Arc RNA (middle) and GFP RNA (right). Insets show fully formed capsids captured under all conditions. Scale bar: 50 nm.

**Fig. S3:**
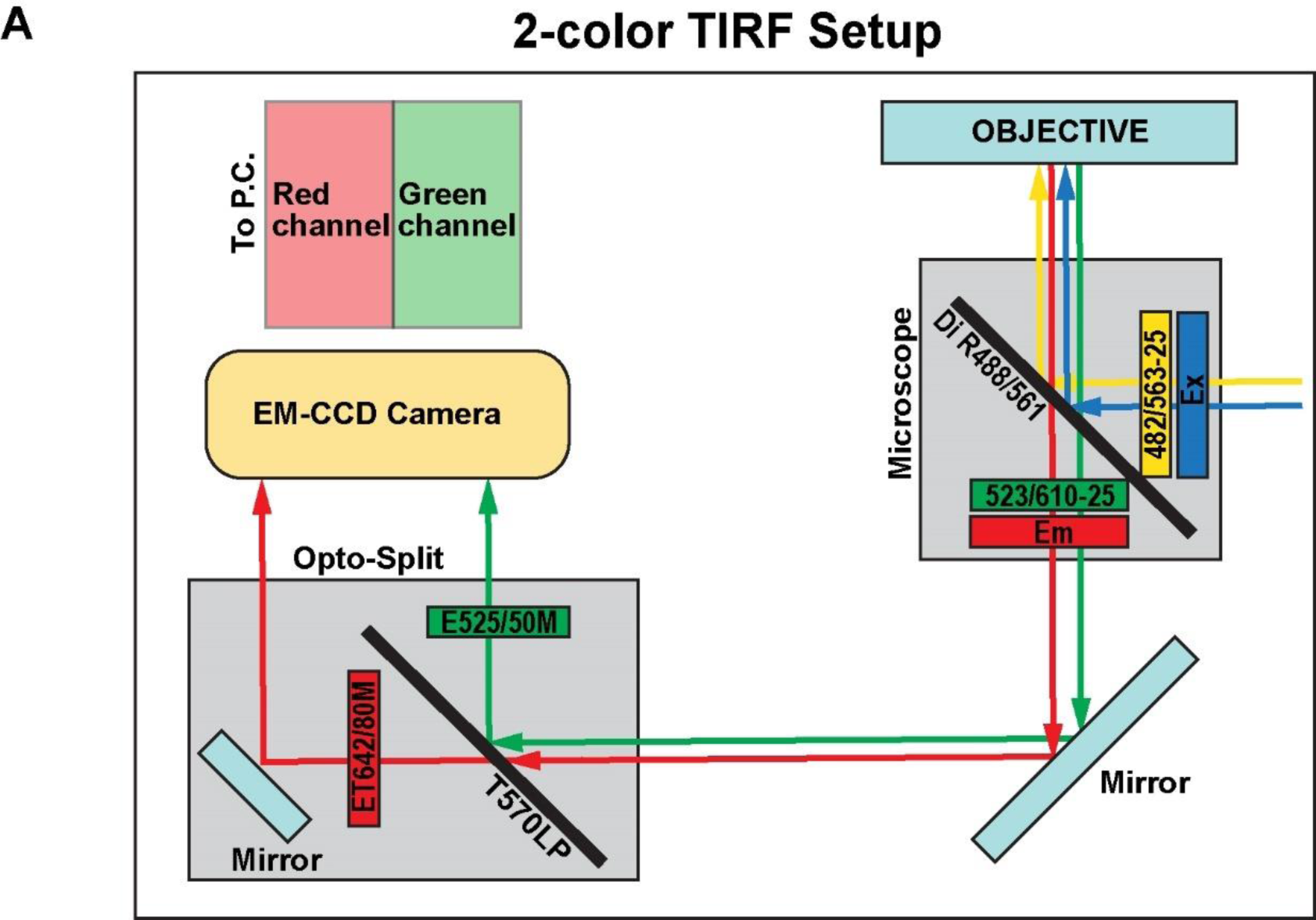
A) Light path for the Opto-Split II setup for 2-color HILO microscopy. The incoming light is split into specific wavelengths and collected separately on the EM-CCD camera for simultaneous tracking of two fluorophores.

**Fig. S4:**
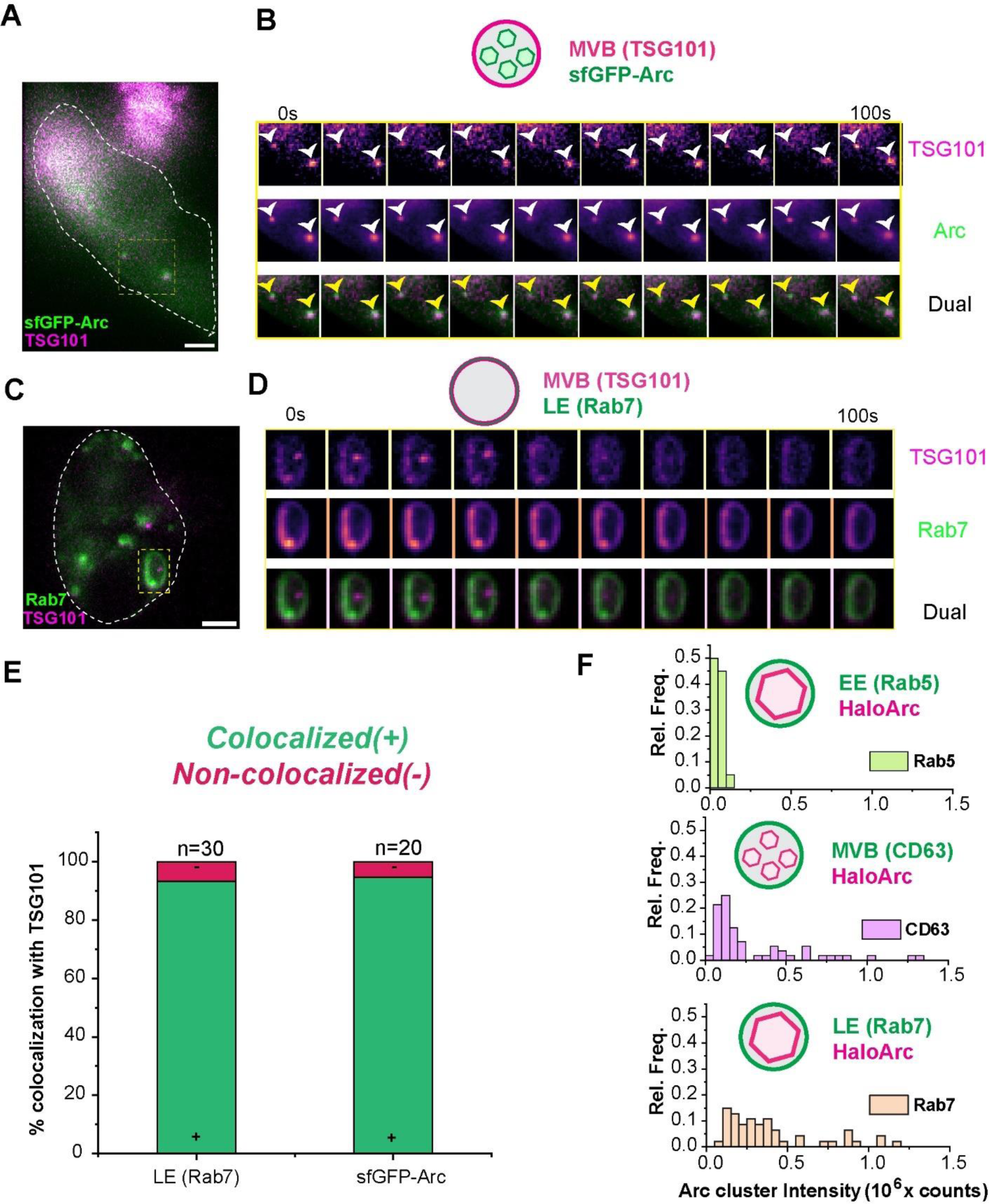
A) Representative image of a HEK-293T cell co-expressing sfGFPArc (green) and TSG101-mCherry (magenta) showing Arc clusters reside inside MVBs vesicles. B) Montage depiction showing images from the ROI shown in A. Warmer colors indicate higher intensities. Arrowheads point Arc clusters. C) A) Representative image of a HEK-293T cell co-expressing Rab7-GFP (green) and TSG101-mCherry (magenta) showing MVB and late endosome markers overlap. D) Montage depiction showing an enlarged vesicle from the ROI shown in C. Warmer colors indicate higher intensities. E) Counting analysis for colocalization (green is + and red is -) with TSG101 with Rab7 (n= 30 vesicles) and TSG101 colocalization with sfGFP Arc cluster (n=20 clusters). F) F.I. distribution of HaloArc clusters residing inside early endosome (top, green), MVB (middle, pink) and late endosome (bottom, orange). Scale bar: 5 µm.

**Fig. S5:**
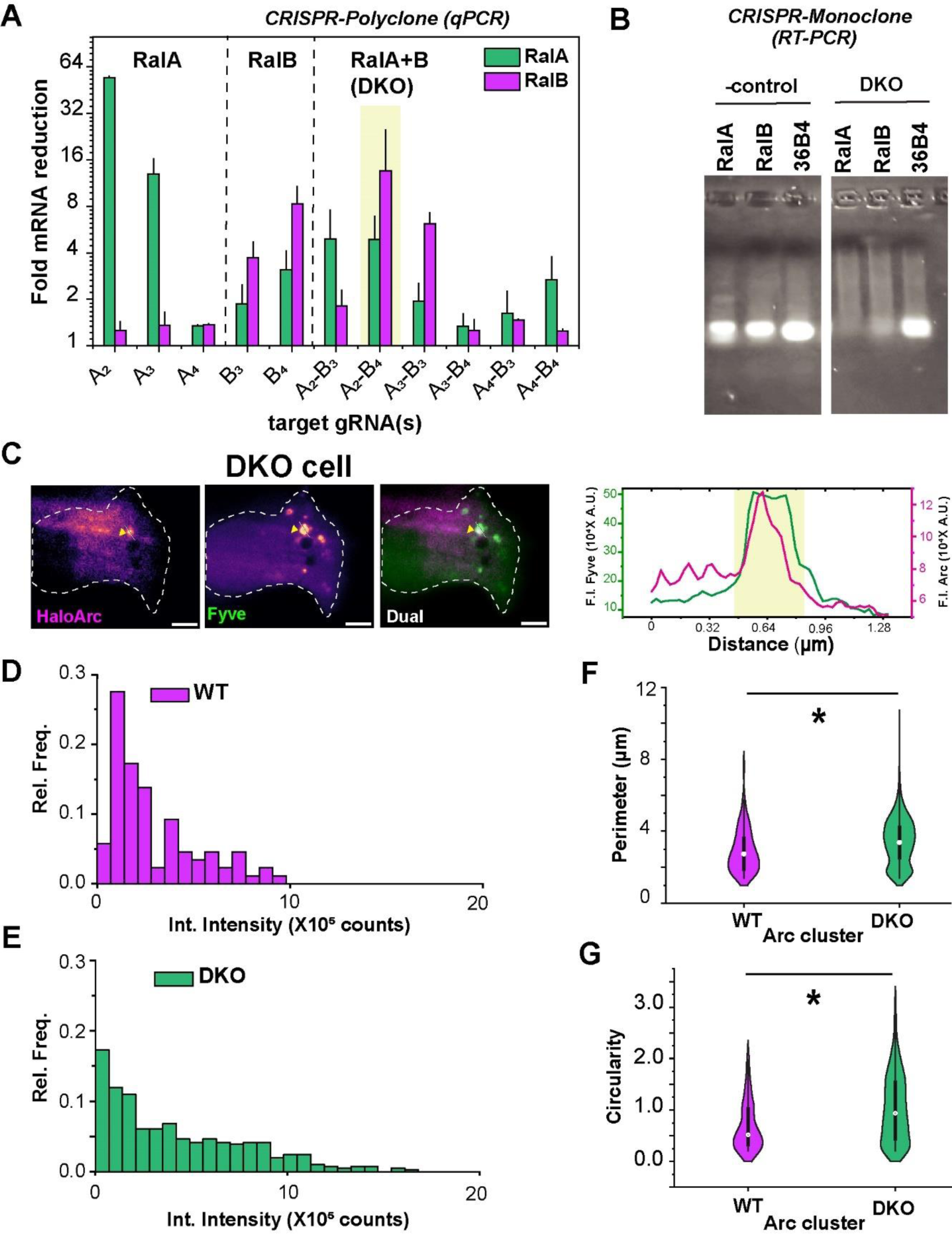
A) Relative fold reduction of RalA and RalB mRNA in polyclonal HEK-293T cells treated with different gRNA combinations for CRISPR-Cas9 knockout using qPCR. Yellow inset highlights that gRNA pair ‘A2-B4’ showed a significant reduction in both RalA and RalB expression. B) RT-PCR gel showing monoclonal colony selected showing complete knockout of both RalA and RalB (right). Endogenous levels of all genes are shown in WT cells (left). 36B4 was used as an internal control for both A and B. C) Representative image of a DKO HEK-293T cell co-expressing HaloArc and 2×FYVE-GFP showing Arc clusters reside inside PI3P(+) vesicles even after the knockout (left). Warmer colors indicate higher intensities. Intensity profile across the line (right). Yellow inset shows overlapping FYVE+ vesicle and Arc cluster. Scale bar: 5 µm. D) F.I. histogram of HaloArc clusters in transfected WT HEK-293T cells. E) Integrated F.I. histogram of HaloArc clusters in transfected DKO HEK-293T cells. F-G) Comparison of perimeter (F) and circularity (G) of HaloArc clusters analyzed in both WT and DKO cells. All data sets are compared using the Mann-Whitney U’s test; *: p-value <0.05.

**Fig. S6:**
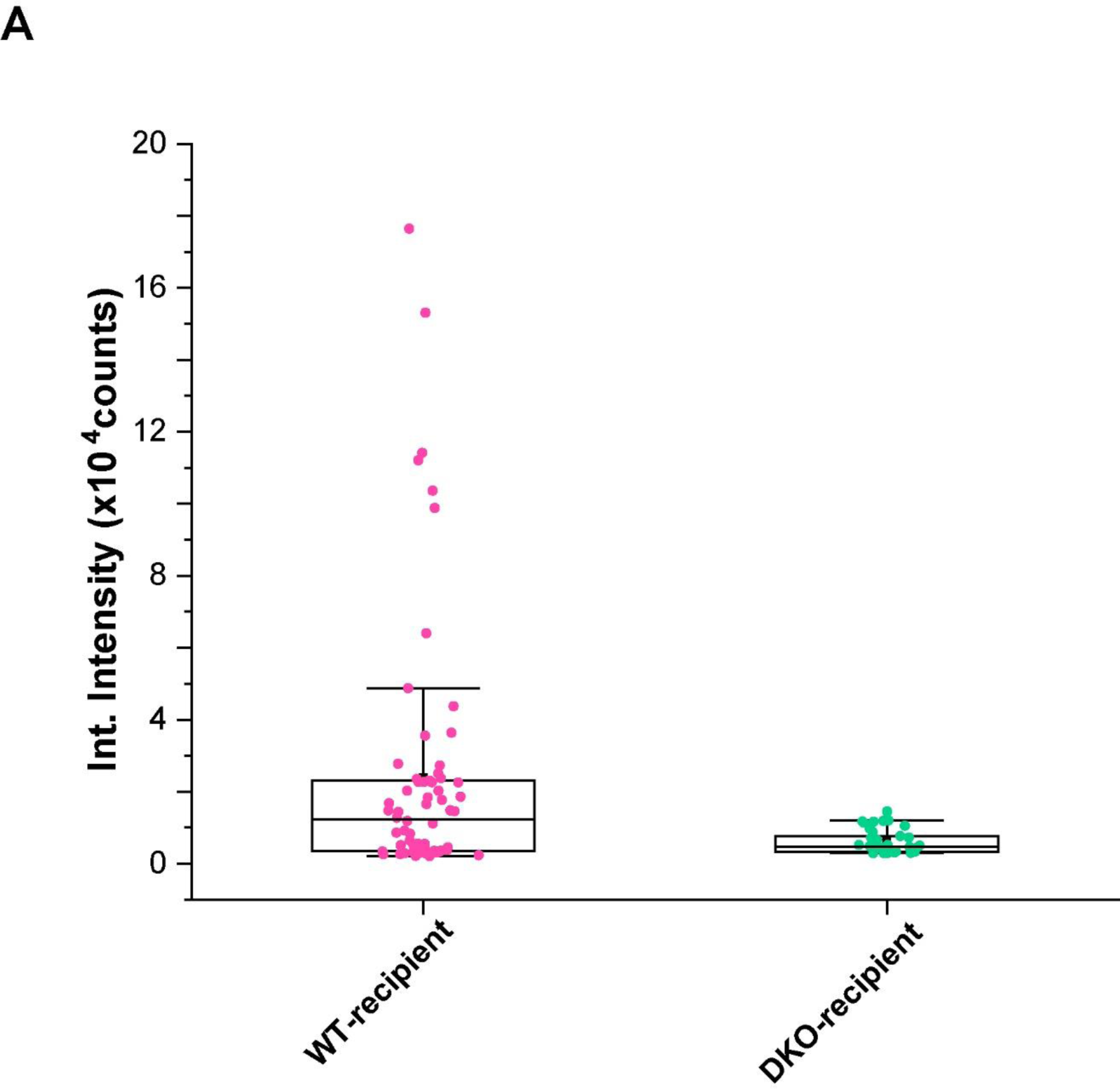
A) Box plot showing intensities of recipient cells positive for HaloArc after media transfer from HaloArc expressing donor cells (left, WT; right, DKO). Recipient cells receiving media from HaloArc-expressing DKO cells show minimal expression of HaloArc.

**Table S1:** Plasmids used in this work.

| Name | Source |
| --- | --- |
| pCS2+-HaloTag-mArc-UTRs | This study |
| pCS2+-sfGFP-mArc-UTRs | This study |
| pCS2+-HaloTag | This study |
| pCS2+-sfGFP | This study |
| pEGFP-hRab5 (Kan) | Zhang Lab, UIUC |
| pEGFP-hRab7 (Kan) | Zhang Lab, UIUC |
| pMEH371 CD63 HA (amp) | Addgene #168220 |
| pCS2+-sfGFP-hCD63 (Amp) | This study (Constructed from Addgene #168220) |
| pEF.6-hTSG101-mCherry (Amp) | Addgene #38318 |
| pEGFP-2xFYVE (Kan) | Addgene #140047 |
| pEGFP-hMTMR1 (Kan) | This study (Constructed from Addgene #100723) |
| pEGFP-hINPP5E (Kan) | This study (Constructed from Addgene #171948) |
| pGEX-6p1-GST-rArcFL (Amp) | Prof. Jason Shepherd, Univ. of Utah |
| pLJM1-Halo-mArc-UTRs (Amp) | This study |
| pCS2+-SnapTag-mArc-UTRs (Amp) | This study (Constructed from Addgene #101135) |

**Table S2:**
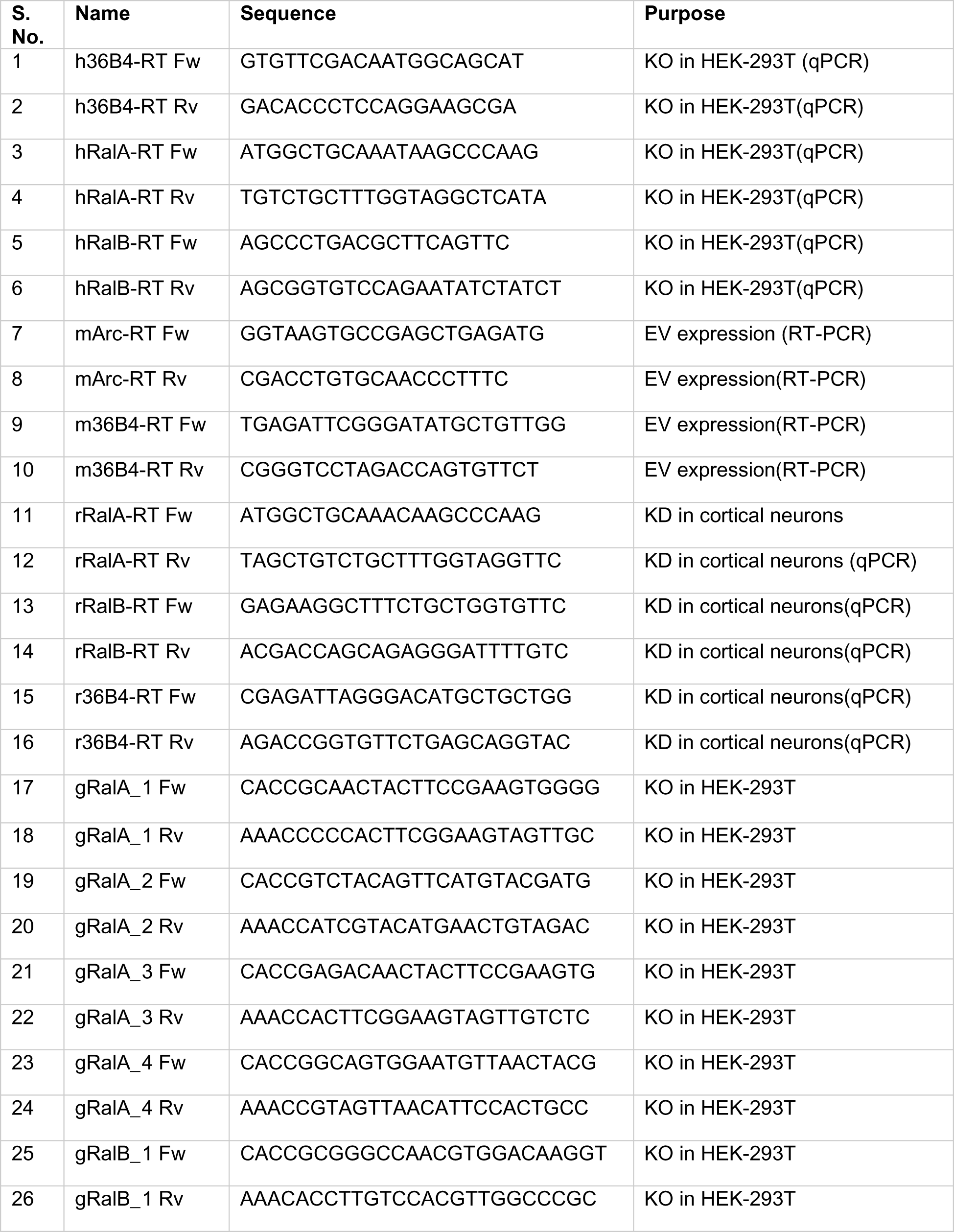
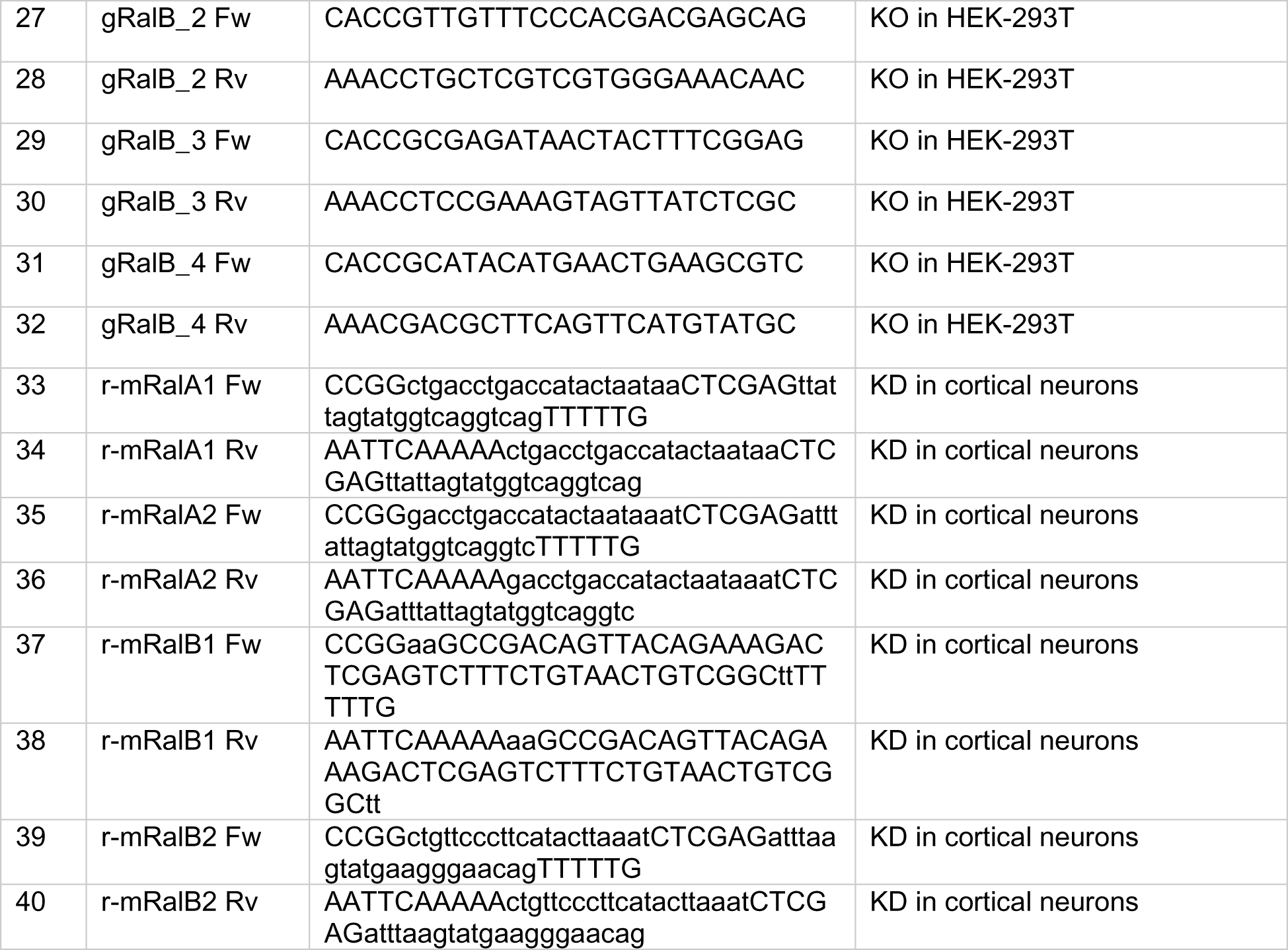
Primers used in this work.

**Table S3:** Reagents used in this work.

| S. No. | Name | Brand | Identifier |
| --- | --- | --- | --- |
| **<u>vectors</u>** |  |  |  |
| 1 | pCS2+ vector | NovoPro | V011726 |
| 2 | pLenti-CRISPRv2 vector | Addgene | 52961 |
| 3 | pLKO.1 TRC cloning vector | Addgene | 10878 |
| 4 | pLJM1 vector | Addgene | 91980 |
| **<u>enzymes/Master-mixes</u>** |  |  |  |
| 5 | FD-EcoRI | Thermo Fisher Scientific | FD0274 |
| 6 | FD-NotI | Thermo Fisher Scientific | FD0593 |
| 7 | FD-PacI | Thermo Fisher Scientific | FD2204 |
| 8 | FD-BcuI | Thermo Fisher Scientific | FD1254 |
| 9 | FD-BshTI | Thermo Fisher Scientific | FD1464 |
| 10 | T4-polynucleotide kinase (T4-PNK) | NEB | M0201 |
| 11 | FD-EspIII | Thermo Fisher Scientific | FD0454 |
| 12 | Shrimp Alkaline phosphatase (rSAP) | NEB | M0371 |
| 13 | Infusion cloning master mix | Takara Bio | 638944 |
| 14 | Reverse Transcriptase | NEB | M0253 |
| 15 | SYBR green master mix | Applied Biosystems | A25742 |
| 16 | Go-Taq green master mix | Promega | M7122 |
| 17 | RNase | Thermo Fisher Scientific | EN0531 |
| 18 | PreScission Protease | Cytiva | 27-0843-01 |
| **<u>Cell culture reagents</u>** |  |  |  |
| 19 | Dulbecco's Modified Eagle Media | Corning | 10-014-CM |
| 20 | Penicillin-streptomycin solution 100X | Corning | 30-002-CI |
| 21 | Fetal bovine serum | Sigma Aldrich |  |
| 22 | PEI max | PolySciences | 24765 |
| 23 | Dulbecco's Modified Phosphate Buffered Saline | Corning | 20-030-CV |
| 24 | Phenol red-free DMEM | Gibco | 31053028 |
| 25 | Trypsin | Life Tech | 15050065 |
| **<u>Antibodies</u>** |  |  |  |
| 26 | Anti-HaloTag | Promega | G9211 |
| 27 | Anti-Arc | Synaptic Systems | 156 003 |
| 28 | Anti-GAPDH | Cell Signaling Technologies | 2118L |
| 29 | Anti-MAP2 | Abcam | Ab5392 |
| 30 | Chicken secondary (Alexa-647) | Abcam | Ab150175 |
| 31 | Mouse secondary HRP | Cell Signaling Technologies | 7076P2 |
| 32 | Rabbit secondary HRP | Cell Signaling Technologies | 7074S |
| **<u>Buffers/Reagents</u>** |  |  |  |
| 33 | RIPA buffer | Millipore Sigma | R0278 |
| 34 | 4X NU-PAGE | Invitrogen | NP0007 |
| 35 | BME | MP Biomedicals | 0219483425 |
| 36 | Bradford Reagent | Thermo Fisher Scientific | 23200 |
| 37 | SDS buffer | Biorad | 1610732 |
| 38 | SDS-PAGE gel | Biorad | 4561043 |
| 39 | Clarity ECL WB substrate | Biorad | 170-5060 |
| 40 | Agarose | Fisher | BP160100 |
| 41 | PVDF membrane | Biorad | 1620177 |
| 42 | Copper 200-mesh grids | Electron Microscopy Sciences | FCF200CU50 |
| 43 | TriZOL reagent | Invitrogen | 15596018 |
| 44 | Chloroform |  |  |
| 45 | Glycogen | Invitrogen | 10814010 |
| 46 | DEPC water | Invitrogen | 462224 |
| 47 | Ethanol | Decon Labs | VN1170 |
| 48 | Isopropanol | VWR | 3032-16 |
| 49 | DTT | Thermo Scientific | 20290 |
| 50 | PMSF | Thermo Scientific | 36978 |
| 51 | LB broth | Fisher Scientific | BP9723 |
| 52 | L-Glutathione | Fisher Scientific | AAJ6216606 |
| 53 | PIP-strip | Echelon Biosciences | P-6001 |
| **<u>Kits</u>** |  |  |  |
| 54 | GeneJet RNA purification kit | Thermo Fisher Scientific | K0731 |
| 55 | Capturem EV isolation mini kit | Takara Bio | 635741 |
| 56 | GeneJet plasmid miniprep | Thermo Fisher Scientific | K0502 |
| 57 | GeneJet Gel extraction kit | Thermo Fisher Scientific | K0691 |
| 58 | MaxiScript SP6 Kit | Invitrogen | AM1340 |
| **<u>Dyes/Fluorophores</u>** |  |  |  |
| 59 | JF-549 halo ligand | Promega | GA1110 |
| 60 | JF-646 halo ligand | Promega | GA1120 |
| 61 | Vybrant DiO cell-labeling solution | Thermo Fisher Scientific | V22886 |
| 62 | AlexaFluor-647 Snap substrate | NEB | S9136S |
| **<u>Antibiotics</u>** |  |  |  |
| 63 | Puromycin | Gold Biotechnology | P-600-100 |
| 64 | Ampicillin | Fisher Scientific | bp176025 |
| 65 | Kanamycin | Fisher Scientific | bp9065 |

**Movie S1 (separate file).** Related to Fig. 2A. Representative HILO imaging of a cell expressing HaloArc (warmer colors indicate higher intensities).

**Movie S2 (separate file).** Related to Fig. 2B. Representative HILO imaging of a cell expressing HaloTag(warmer colors indicate higher intensities)

**Movie S3A (separate file).** Related to Fig. 4C. Representative HILO imaging of a WT HEK-293T cell co-expressing HaloArc (magenta) and 2×FYVE-GFP (green) showing colocalization of Arc clusters with PI3P

**Movie 3B (separate file).** Related to Fig. S3B. Representative HILO imaging of a DKO HEK-293T cell co-expressing HaloArc (magenta) and 2×FYVE-GFP (green) showing colocalization of Arc clusters with PI3P

**Movie S4A (separate file).** Related to Fig. 5A. Representative HILO imaging of a cell co-expressing HaloArc (magenta) and Rab5-GFP (green) showing localization of Arc clusters with early endosome

**Movie S4B (separate file).** Related to Fig. 5B. Representative HILO imaging of a cell co-expressing HaloArc (magenta) and CD63-sfGFP (green) showing localization of Arc clusters with MVBs

**Movie S4C (separate file).** Related to Fig. 5C. Representative HILO imaging of a cell co-expressing HaloArc (magenta) and Rab7-GFP (green) showing localization of Arc clusters with late endosomes and MVBs

**Movie S4D (separate file).** Related to Fig. S4A. Representative HILO imaging of a cell co-expressing MVB marker TSG101-mCherry (magenta) and Rab7-GFP (green) showing late endosomes and MVBs can have common markers.

**Movie S4E (separate file).** Related to Fig. S4B. Representative HILO imaging of a cell co-expressing sfGFP-Arc (green) and TSG101-mCherry (magenta) showing Arc clusters also localize with ESCRT protein TSG101.

**Dataset S1 (separate file).** Diffusion Coefficient values for HaloArc and HaloTag tracked particles using the Co-variance estimator method.

**Dataset S2 (separate file).** Step size and fluorescence intensities of Arc clusters colocalizing with EE, MVB and LE.

**Dataset S3 (separate file).** WT and DKO Arc cluster analysis data for fluorescence intensity, cluster number per cell, integrated intensity, perimeter and circularity.

**Software S1 (separate file).** Custom-written script for Diffusion Coefficient analysis performed in the paper using the CVE method. https://github.com/kritikamehta1794/Diffusion-coefficient_CVE.git

